# Cell size scaling laws: a unified theory

**DOI:** 10.1101/2022.08.01.502021

**Authors:** Romain Rollin, Jean-François Joanny, Pierre Sens

## Abstract

The dimensions and compositions of cells are tightly regulated by active processes. This exquisite control is embodied in the robust scaling laws relating cell size, dry mass, and nuclear size. Despite accumulating experimental evidence, a unified theoretical framework is still lacking. Here, we show that these laws and their breakdown can be explained quantitatively by three simple, yet generic, physical constraints defining altogether the Pump and Leak model (PLM). Based on estimations, we clearly map the PLM coarse-grained parameters with the dominant cellular events they stem from. We propose that dry mass density homeostasis arises from the scaling between proteins and small osmolytes, mainly amino-acids and ions. Our theory predicts this scaling to naturally fail, both at senescence when DNA and RNAs are saturated by RNA polymerases and ribosomes respectively, and at mitotic entry due to the counterion release following histone tail modifications. We further show that nuclear scaling result from osmotic balance at the nuclear envelope (NE) and a large pool of metabolites, which dilutes chromatin counterions that do not scale during growth.

## I. INTRODUCTION

Although cell size varies dramatically between cell types, during the cell cycle and depends on various external stresses [1], each cell type often shows small volumetric variance. This tight control reflects the importance of size in monitoring cell function. It is often associated to generic linear scaling relations between cell volume, cell dry mass and the volume of the nucleus ([2], [3], [4]). These scaling laws have fascinated biologists for more than a century [5] [6], because of the inherent biological complexity and their ubiquity both in yeasts, bacteria and mammals, hence raising the question of the underlying physical laws.

Although robust, these scaling relations do break down in a host of pathologies. The nuclear-to-cytoplasmic (NC) ratio (also called karyoplasmic ratio) has long been used by pathologists to diagnose and stage cancers ([7],[8],[9]). Similarly, senescent cells such as fibroblasts are known to be swollen and their dry mass diluted [10], a feature suspected to be of fundamental biological importance since it could represent a determinant of stem cell potential during ageing [11].

Paradoxically, there is still no unified understanding of these scaling laws and of the reasons of their breakdown in diseases. This is in part due to the experimental difficulty to perform accurate volume and dry mass measurements ([12],[13],[14]). Many methods were developed in the past decades but they sometimes lead to contradictory observations highlighting the need of comparing and benchmarking each method ([15],[16]).

Moreover, extensive experimental investigations have identified a plethora of biological features influencing these scalings but comparatively fewer theoretical studies have precisely addressed them, leaving many experimental data unrelated and unexplained. Several phenomenological theories have emerged to understand individual observations, but they are still debated among biologists. The “nucleoskeletal theory” emphasizes the role of the DNA content in controling the NC ratio, based on the idea that ploidy dictates cell and nuclear sizes since tetraploid cells tend to be larger than their diploid homologs [4]. Other experiments suggest that genome size is not the only determining factor: indeed it would not explain why cells from different tissues, having the same amount of DNA, have different sizes. Instead, it has been shown that nuclear size depends on cytoplasmic content, nucleo-cytoplasmic transport, transcription, RNA processing and mechanics of nuclear envelope structural elements such as Lamina [3].

In parallel, theoretical models, based on non-equilibrium thermodynamics, were developed ([17],[18],[19]), often based on the “Pump-and-Leak” principle ([1],[16],[20]). Charged impermeant molecules in cells create an imbalance of osmotic pressure at the origin of the so-called Donnan effect [21]. Cells have two main ways to counteract the osmotic imbalance. They can adapt to sustain a high hydrostatic pressure difference as plants do by building cellulose walls. Or, as done by mammalian cells, they can use ion pumps to actively decrease the number of ions inside the cells, thus decreasing the osmotic pressure difference across the cell membrane and therefore impeding water penetration. However, due to the large number of parameters of these models, we still have a poor understanding of the correspondence between biological factors and physical parameters of the model.

In this paper, we bridge the gap between phenomenological and physical approaches by building a minimal framework based on a nested PLM to understand the cell size scaling laws as well as their breakdown. Performing order of magnitude estimates, we precisely map the coarse-grained parameter of a simplified version of the PLM to the main microscopic biological events. We find that the dry mass of the cell is dominated by the contribution of the proteins, while the cell volume is mostly fixed by the contribution to the osmotic pressure of small osmolytes, such as aminoacids and ions. The maintenance of a homeostatic cell density during growth is then due to a linear scaling relation between protein and small osmolyte numbers. Combining simplified models of gene transcription and translation and of amino-acid biosynthesis to the PLM, we show that the linear scaling relation between protein and small osmolyte numbers is obtained in the exponential growth regime of the cell by virtue of the enzymatic control of amino-acid production.

On the other hand, the absence of linear scaling relation between protein and small osmolyte numbers is at the root of the breakdown of density homeostasis. We show that this is the case both at senescence and at mitotic entry due to two distinct physical phenomena. At senescence, cells cannot divide properly. Our theory then predicts that DNA and RNAs become saturated by RNA polymerases (RNAPs) and ribosomes respectively, leading to a change of the growth regime: the protein number saturates while the amino-acid number increases linearly with time, resulting in the experimentally observed dry mass dilution. This prediction is quantitatively tested using published data of growing yeast cells prevented from dividing [10]. At mitotic entry, chromatin rearrangements, such as histone tail modifications, induce a counterion release inside the cell, resulting in an influx of water and dry mass dilution in order to maintain the osmotic balance at the cell membrane.

Finally, to further illustrate the generality of our model, we show that the linear scaling of nucleus size with cell size originates from the same physical effects. Using a nested PLM for the cell and the nucleus, we show that nuclear scaling requires osmotic balance at the nuclear envelope. The osmotic balance is explained by the nonlinear osmotic response of mammalian nuclei, that we attribute to the presence of folds at the surface of many nuclei [22], which in turn buffer the NE tension and enforce scaling. Nonetheless, the condition on osmotic balance appears to be insufficient to explain the robustness of the NC ratio during growth. Counter-intuitively, metabolites, though permeable to the NE, are predicted to play an essential role in the NC ratio. Their high concentrations in cells, a conserved feature throughout living cells, is shown to dilute the chromatin counterions which do not scale during growth; thereby allowing the scaling of nuclear size with cell size both at the population level and during individual cell growth.

## II. RESULTS

### A. Pump and leak model

Our theoretical approach to understand the various scaling laws associated to cell size is based on the the Pump and Leak model (PLM) ([23] and Figure1.A). The PLM is a coarse grained model emphasizing the role of mechanical and osmotic balance. The osmotic balance involves two types of osmolytes, impermeant molecules such as proteins and metabolites, which cannot diffuse through the cell membrane, and ions, which cross the cell membrane and for which at steady state, the incoming flux into the cell must equal the outgoing flux. For simplicity, we restrict ourselves to a two-ions PLM where only cations are pumped outward of the cell. We justify in the Discussion why this minimal choice is appropriate for the purpose of this paper. Within this framework, three fundamental equations determine the cell volume. (1) Electroneutrality: the laws of electrostatic ensure that in any aqueous solution such as the cytoplasm, the solution is neutral at length scales larger than the Debye screening length i.e. the electrostatic charge of any volume larger than the screening length vanishes. In physiological conditions, the screening length is typically on the nanometric scale. Therefore, the mean charge density of the cell vanishes in our coarse-grained description (Eq.S.1) (2) Osmotic balance: balance of the chemical potential of water inside and outside the cell; the timescale to reach the equilibrium of water is of the order of tens to hundreds of milliseconds after a perturbation [16],[24]. (3) Balance of ionic fluxes: the typical timescales of ion relaxation observed during a cell regulatory volume response after an osmotic shock are of the order of a few minutes [16], [25]. Together, this means that our quasi-static theory is designed to study cell size variations on timescales larger than a few minutes. Mathematically, the three equations read (see Appendix I for the full derivations of these equations) :

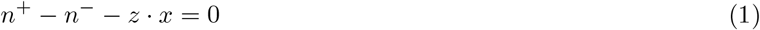

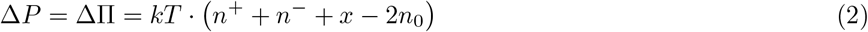

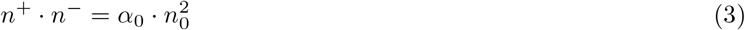

where, *n*^+^, *n*^−^, *n*_0_ are respectively the cationic and anionic concentrations inside and outside the cell. The external ionic concentrations are assumed to be identical for cations and anions in order to enforce electroneutrality since the concentrations of non-permeant molecules in the external medium are typically much lower than their ionic counterparts [24]. The cell is modelled as a compartment of total volume *V* divided between an excluded volume occupied by the dry mass *R* and a wet volume. The cell contains ions and impermeant molecules such as proteins, RNA, free amino-acids and other metabolites. The number *X*, respectively the concentration 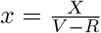, of these impermeant molecules may vary with time due to several complex biochemical processes such as transcription, translation, plasma membrane transport, and degradation pathways. The average negative charge −*z* of these trapped molecules induces a Donnan potential difference *U*_*c*_ across the cell membrane. The Donnan equilibrium contributes to the creation of a positive difference of osmotic pressure ΔΠ that inflates the cell. Cells have two main ways to counteract this inward water flux. They can either build a cortex stiff enough to prevent the associated volume increase, as done by plant cells. This results in the appearance of a hydrostatic pressure difference Δ*P* between the cell and the external medium. Or they can pump ions outside the cell to decrease the internal osmotic pressure, a strategy used by mammalian cells. We introduce a pumping flux of cations *p*. Cations can also passively diffuse through the plasma membrane via ion channels with a conductivity *g*^+^. In Eq.3, the pumping efficiency is measured by the dimensionless number 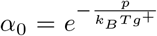 where *T* is the temperature and *k*_*B*_ the Boltzmann constant. The pumping efficiency varies between 0 in the limit of “infinite pumping” and 1 when no pumping occurs (see Appendix I for an explanation on the origin of this parameter).

**FIG. 1.**
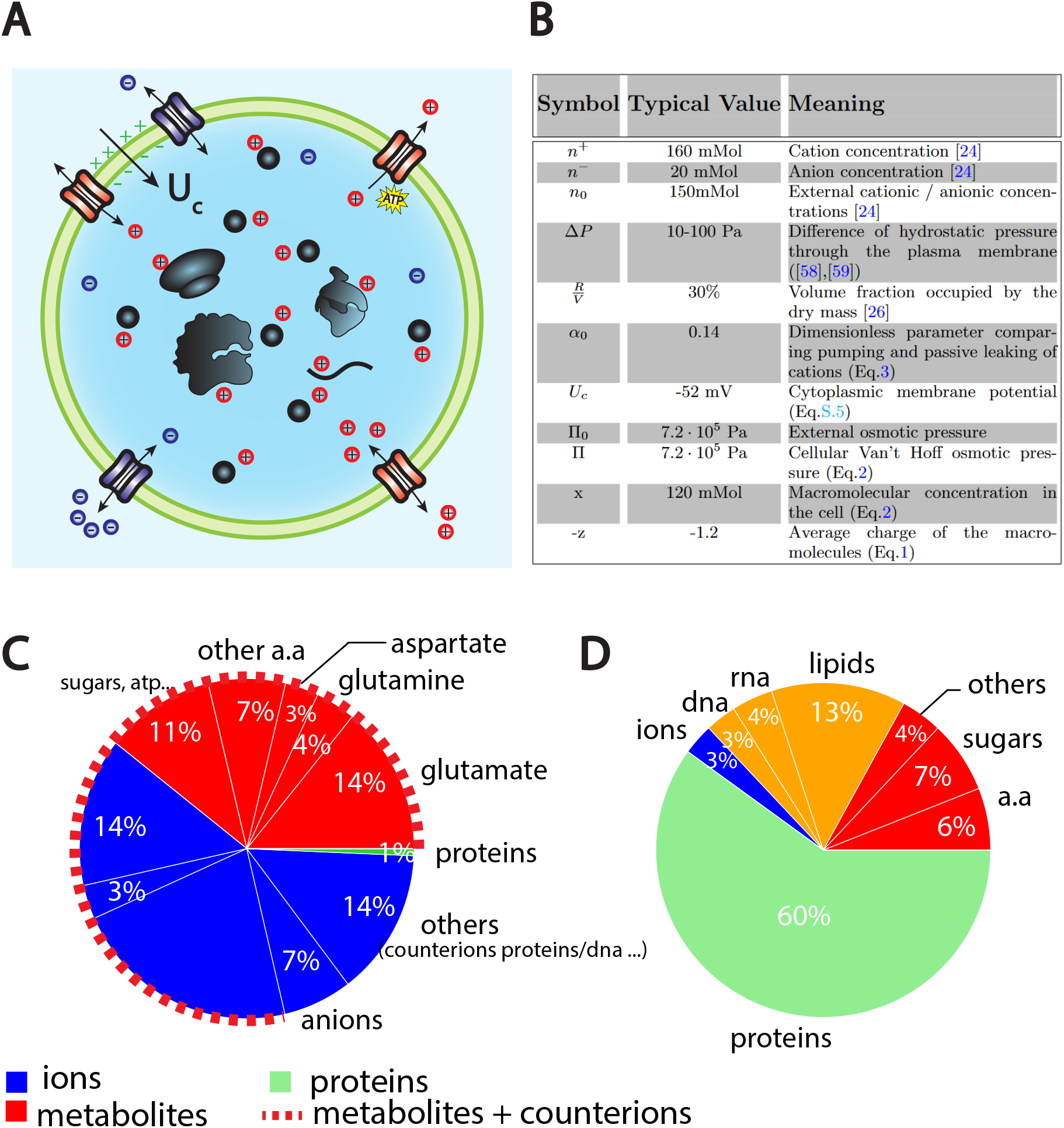
The PLM. (A) Schematic of the PLM. Species in black are impermeant molecules such as proteins, mRNAs and metabolites (black circles). In average, those molecules are negatively charged and thus carry positive counterions (red species) to ensure electroneutrality. Ions can freely cross the plasma membrane through channels. Their concentrations in the cell result from the balance of three fluxes: the electrical conduction, the entropic diffusion, and pumping. In the model, only cations are pumped out of the cell to model the Na/K pump but this assumption is not critical (see Discussion III C and Appendix II C) (B) Estimation of the coarse grained PLM parameters for a typical Mammalian cell. (C) Fraction of volume and (D) of the dry mass occupied by the constituents of a mammalian cell (see Appendix III and [26]). Note that most of the number of impermeant molecules (X) are accounted for by metabolites (mainly amino-acids and glutamate).

### B. Volume and dry mass scaling

Although proposed more than 60 years ago [23] and studied in depth by mathematicians [18], and physicists [27], little effort has been done to precisely map the coarse-grained parameters of the PLM to microscopic parameters. We adopt here the complementary strategy and calculate orders of magnitude in order to simplify the model as much as possible, only keeping the leading order terms. We summarize in Figure.1.B the values of the PLM parameters that we estimated for a “typical” mammalian cell. Three main conclusions can be drawn: (1) Pumping is important, as indicated by the low value of the pumping efficiency *α*_0_ ∼ 0.14. Analytical solutions presented in the main text will thus be given in the “infinite pumping” limit, i.e., *α*_0_ ∼ 0, corresponding to the scenario where the only ions present in the cell are the counterions of the impermeant molecules. Though not strictly correct, this approximation gives a reasonable error of the order 10% on the determination of the volume, due to the typical small concentration of free anions in cells Fig.1.B. This error is comparable to the typical volumetric measurement errors found in the literature. (2) Osmotic pressure is balanced at the plasma membrane of a mammalian cell, since hydrostatic and osmotic pressures differ by at least three orders of magnitude. This result implies that even though the pressure difference Δ*P* plays a significant role in shaping the cell, it plays a negligible role in fixing the volume (see Eq.S.12 for justification). (3) The cellular density of impermeant species is high, *x* ∼ 120mMol, comparable with the external ionic density *n*_0_.

In this limit, the volume of the cell hence reads (the complete expression is given in Appendix II) :

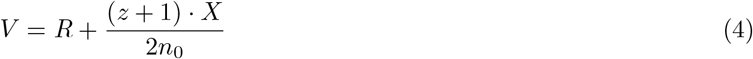

The wet volume of the cell is thus slaved to the number of impermeant molecules that the cell contains. While this conclusion is widely acknowledged, the question is to precisely decipher which molecules are accounted for by the number X. We first estimate the relative contributions of the cellular free osmolytes to the volume of the cell and then, compute their relative contributions to the dry mass of the cell. We provide a graphical summary of our orders of magnitudes in Fig.1.C and D as well as the full detail of their derivations in Appendix.III. The conclusion is twofold. Metabolites and their counterions account for most of the wet volume of the cell, 78% of the wet volume against 1% for proteins. On the other hand, proteins account for most of the dry mass of mammalian cells, accounting for 60% of the cellular dry mass against 17% for metabolites.

We further note that metabolites are mainly amino-acids and in particular three of them, glutamate, glutamine and aspartate accounting for 73% of the metabolites [28]. It is important to note that the relative proportion of free amino-acids in the cell does not follow their relative proportion in the composition of proteins. For instance, glutamate represents 50% of the free amino-acid pool while its relative appearance in proteins is only 6% [29]. This is evidence that some amino-acids have other roles than building up proteins. In particular, we demonstrate throughout this paper their crucial role on cell size and its related scaling laws.

These conclusions may appear surprising due to the broadly reported linear scaling between volume (metabolites) and dry mass (proteins), hence enforcing a constant dry mass density *ρ* during growth. Many theoretical papers have assumed a priori a linear phenomenological relation between volume and protein number in order to study cell size [30],[31],[32]. Our results instead emphasize that the proportionality is indirect, only arising from the scaling between amino-acid and protein numbers. The dry mass density reads (to lowest order):

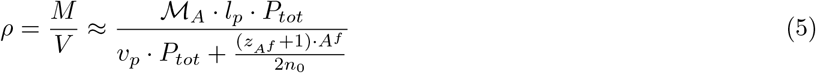

where, ℳ_*a*_, *z*_*Af*_ and *A*^*f*^ are respectively the average mass, charge and number of amino-acids; *l*_*p*_, *v*_*p*_ and *P*_*tot*_, the average length, excluded volume and number of proteins. Note that density homeostasis is naturally achieved in the growth regime where *A*^*f*^ is proportional to *P*_*tot*_.

### C. Model of stochastic gene expression and translation

To further understand the link between amino-acid and protein numbers we build upon a recent model of stochastic gene expression and translation ([30] and Fig.2.A). The key feature of this model is that it considers different regimes of mRNA production rate 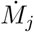 and protein production rate 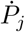 according to the state of saturation of respectively the DNA by RNA polymerases (RNAPs) and mRNAs by ribosomes. For the sake of readability, we call enzymes both ribosomes and RNAPs, their substrates are respectively mRNAs and DNA and their products proteins and mRNAs. The scenario of the model is the following. Initially, the majority of enzymes are bound to their substrates and occupy a small fraction of all possible substrate sites. In this non saturated regime, i.e when the number of enzymes is smaller than a threshold value 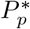 and 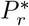 Eq.S.34, the production rates of the products of type j read [30]:

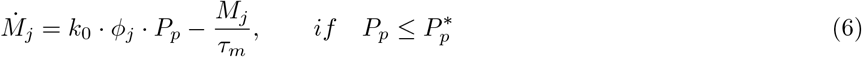

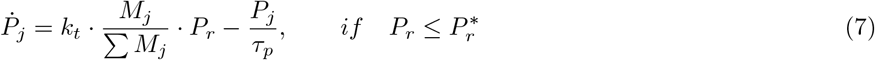

Both production rates have two contributions. (1) a source term characterized by the rates *k*_0_ and *k*_*t*_ at which the enzyme produces the product once it is bound to its substrate, times the average number of enzymes per substrate coding for the product of type j. This number is the fraction of substrates coding for product of type j - that can be identified as a probability of attachment (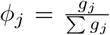 and 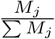, where *g*_*j*_, *M*_*j*_ accounts for the number of genes and mRNAs coding for the product of type j) - multiplied by the total number of enzymes (*P*_*p*_ and *P*_*r*_). (2) A degradation term characterized by the average degradation times *τ*_*m*_ and *τ*_*p*_ of mRNAs and proteins. Note that we added a degradation term for proteins not present in [30], which turns out to be of fundamental importance below. Although these timescales vary significantly between species their ratio remains constant, *τ*_*m*_ being at least one order of magnitude smaller than *τ*_*p*_ in yeast, bacteria and mammalian cells [24]. This justifies a quasistatic approximation, 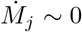 during growth such that the number of mRNAs of type j adjusts instantaneously to the number of RNAPs, in the non saturated regime :

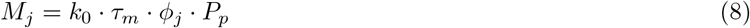

During interphase, the number of enzymes grows, increasingly more enzymes attach to the substrates up to the saturation value due to their finite size. In this regime, we use the same functional form for the production rates only replacing the average number of enzymes per substrate by their saturating values : 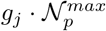 for RNAPs and 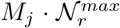 for ribosomes (see Appendix IV and Eq.S.32,S.33); where, 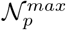 and 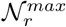 are the average maximal number of RNAPs and ribosomes per mRNAs and genes. Note that the model predicts that the saturation of DNA precedes that of mRNAs, whose number initially increases with the number of RNAPs Eq.8 while the number of genes remains constant. Once DNA is saturated, the number of mRNAs plateaus, leading to their saturation by ribosomes (see Appendix IV and Eq.S.38).

**FIG. 2.**
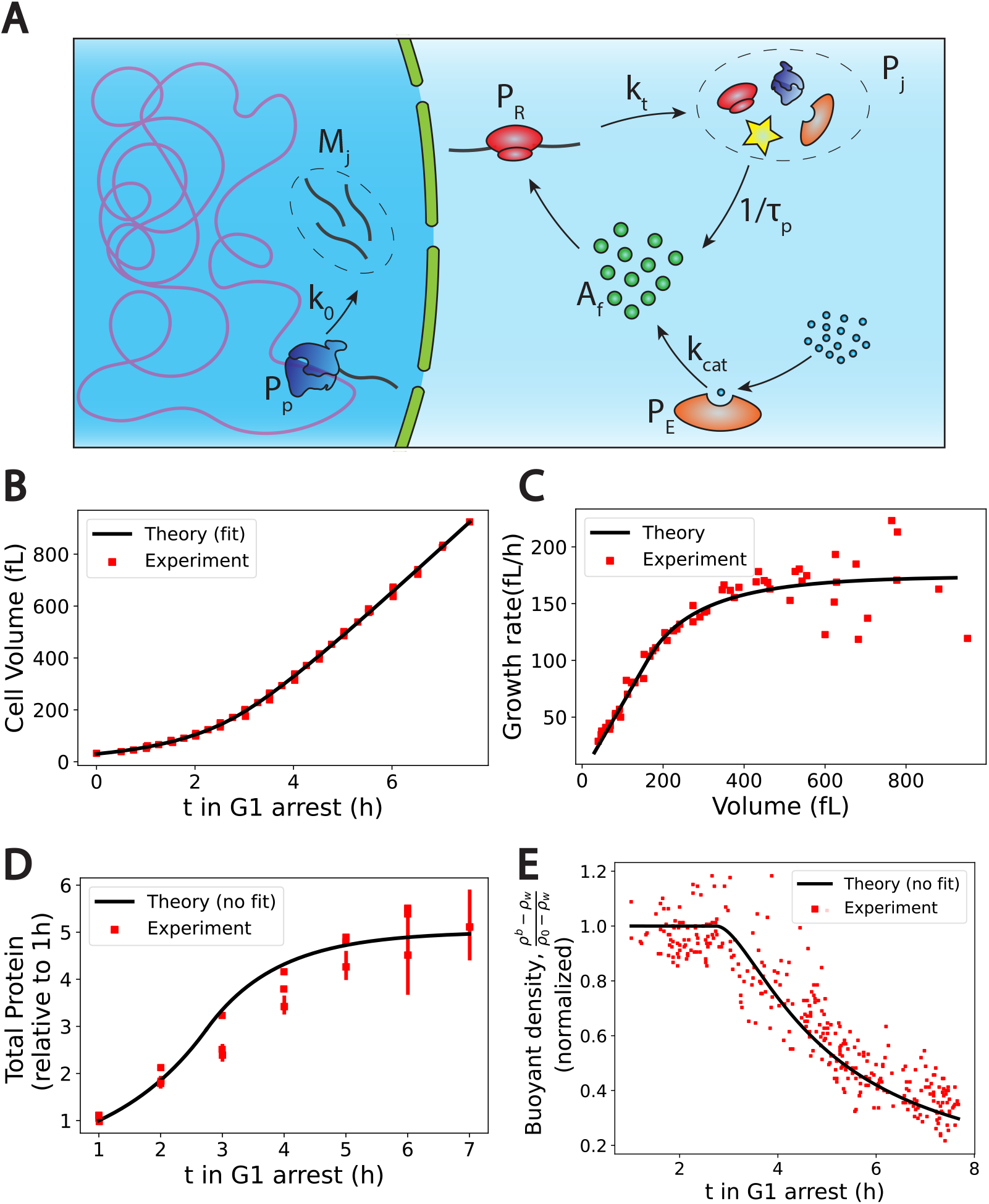
The PLM coupled to a growth model predicts quantitatively dry mass homeostasis and its subsequent dilution at senescence. (A) Schematic of the growth model. RNAPs (*P*_*p*_) transcribe DNA and form mRNAs (*M*_*j*_) at an average rate *k*_0_. mRNAs are then read by ribosomes (*P*_*r*_) to produce proteins (*P*_*j*_) at an average rate *k*_*t*_. Proteins are degraded at an average rate 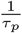 into free amino-acids (*A*_*f*_). Free amino-acids are also synthetized from nutrients (blue circles) at a rate *k*_*cat*_. This reaction is catalyzed by enzymes (*P*_*e*_). (B) to (E) Comparison between theory (black) and experiment (red). Data adapted from [10]. (D) and (E) model predictions without any fitting parameters. The buoyant mass *ρ*_*b*_ density is defined as the total mass of the cell (water and dry mass) over the total volume of the cell.

Our previous analysis has highlighted the fundamental importance of free amino-acids on cell volume regulation Fig.1.C. The production rate of free amino-acids can be related to the number of enzymes catalyzing their biosynthesis, using a linear process by assuming that the nutrients necessary for the synthesis are in excess:

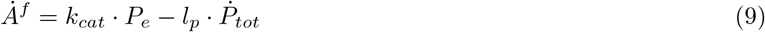

where *k*_*cat*_ is the rate of catalysis and *P*_*e*_ the number of enzymes. The second term represent the consumption of amino-acids to form proteins, with *P*_*tot*_ = ∑ *P*_*j*_. Although Eq.9 is coarse-grained we highlight that, since glutamate and glutamine are the most abundant amino acids in the cell, it could in particular model the production of these specific amino-acids from the Krebs cycle [26]. Note that we also ignored amino-acid transport through the plasma membrane. The rationale behind this choice is twofold. (1) We do not expect any qualitative change when adding this pathway to our model since amino-acid transport is also controlled by proteins. (2) We realized that the amino-acids that actually play a role in controlling the volume, mainly glutamate, glutamine and aspartate, are non-essential amino-acids, hence that can be produced by the cell.

### D. Dry mass scaling and dilution during cell growth

We now combine the PLM, the growth model and the amino-acid biosynthesis model to make predictions on the variation of the dry mass density during interphase. A crucial prediction of the growth model is that as long as mRNAs are not saturated, i.e., 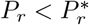, all the protein numbers scale with the number of ribosomes, 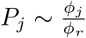 · *P*_*r*_. Moreover, the autocatalytic nature of ribosome formation makes their number grow exponentially Eq.7, i.e 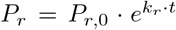; where, 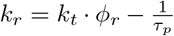 is the effective rate of ribosome formation (and also the rate of volume growth in this regime Eq.S.36). The most important consequence of this exponential growth coupled to the equation modeling amino-acid biosynthesis Eq.9 is that it implies that both amino-acids and total protein content scale with the number of ribosomes ultimately leading to a homeostatic dry mass density independent of time (see Appendix IV):

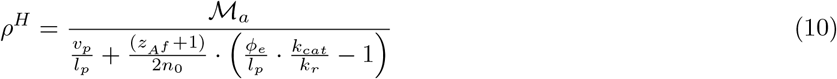

We emphasize that Eq.10 only applies far from its singularity since it was obtained assuming that the volume of the cell is determined by free amino-acids, i.e, 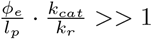.

Not only does our model explain the homeostasis of the dry mass, but it also makes the salient prediction that this homeostasis naturally breaks down if the time spent in the G1 phase is too long. Indeed, after a time 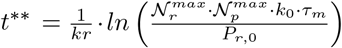 (see AppendixIV), mRNAs become saturated by ribosomes, drastically changing the growth of proteins from an exponential growth to a plateau regime where the number of proteins remains constant. After the time *t*^**^ + *τ*_*p*_, all protein numbers reach their stationary values 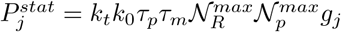. In particular, the enzymes coding for amino-acids also plateau implying the loss of the scaling between free amino-acids and proteins as predicted by Eq.9. The number of amino-acids then increases linearly with time whereas the number of proteins saturates. In this regime, the volume thus grows linearly with time but the dry mass remains constant leading to its dilution and the decrease of the dry mass density (see Appendix IV and Eq.S.42) :

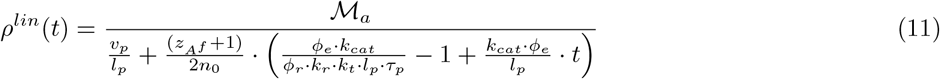

Finally, our model makes other important predictions related to the cell ploidy that we briefly enumerate. First, the cut-off 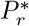 Eq.S.38 at which dilution is predicted to occur depends linearly on the genome copy number ∑*g*_*j*_. Intuitively, mRNAs are saturated only if DNA has previously saturated. At saturation the RNA number is proportional to the genome size. As a consequence, the volume 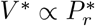 at which dilution occurs scales with the ploidy of the cell, a tetraploid cell is predicted to be diluted at twice the volume of its haploid homolog. On the other hand, by virtue of the exponential growth, the time *t*^**^ Eq.S.39 at which the saturation occurs only depends logarithmically on the number of gene copies making the ploidy dependence much less pronounced timewise. Second, the growth rate in the linear regime scales with the ploidy of the cell, as opposed to the growth rate in the exponential regime. Indeed, in the saturated regime, the growth rate scales as 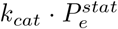 (see Appendix IV and Eq.S.40), where 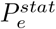 is the number of enzymes catalyzing the reaction of amino-acids biosynthesis after their numbers have reached their stationary values, while in the exponential regime, the growth rate 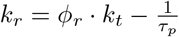 scales with the fraction of genes coding for ribosomes *ϕ*_*r*_, which is independent of the ploidy.

### E. Comparison to existing data

Our main prediction, namely that the cell is diluted after the end of the exponential growth, is reminiscent of the intracellular dilution at senescence recently reported in fibroblasts, yeast cells and more recently suspected in aged hematopoietic stem cells [10],[11]. Here we quantitatively confront our theory to the data of [10], where the volume, the dry mass and the protein number were recorded during the growth of yeast cells that were prevented from both dividing and replicating their genome. Though our theory was originally designed for mammalian cells, it can easily be translated to cells with a cell wall provided that the hydrostatic pressure difference across the wall Δ*P* is maintained during growth by progressive incorporation of cell wall components (see Appendix II). Indeed, our conclusions rely on the fact that the cell volume is primarily controlled by small osmolytes whose concentration in the cell dominates the osmotic pressure, a feature observed to be valid across cell types [28].

We first check the qualitative agreement between our predictions and the experiments. Two clear and very distinct growth regimes are evident in non-dividing yeast cells; an initial exponential growth followed by a linear growth Fig.2.C. The occurrence of linear or exponential growth has been the object of intense debate. We think that the ambiguity comes from the fact that cells often divide too fast for the exponential regime to be properly identified. Our results suggest that the fact that cell division occurs in the exponential regime is essential to prevent cells from being diluted. Our theory also predicts that as long as protein number is constant the volume must grow linearly Eq.9,S.42. This is precisely what is observed in the experiments: cells treated with rapamycin exhibits both a constant protein content and a linear volume increase during the whole growth (see Fig.S6.F in [10]). Finally, our predictions on the relationship between ploidy and dilution are in very good agreement with experiments as well. Indeed, while the typical time to reach the linear growth regime - of the order of 3 hours - seems independent of the ploidy of the cell, the volume at which dilution occurs is doubled (see Fig.S7.A in [10]). Moreover, the growth rate during the linear regime scales with ploidy, as the haploid cells growth rate is of order 129 fL*/*h against 270 fL*/*h for their diploid counterparts [10].

Encouraged by these qualitative correlations, we further designed a scheme to test our theory more quantitatively. Although our theory has a number of adjustable parameters, many of them can be combined or determined self-consistently as shown in Appendix IV D). We end up fitting four parameters, namely *τ*_*p*_, *t*^**^, *k*_*r*_ and the initial cell volume *v*_1_, using the cell volume data Fig.2.B. We detail in the Appendix IV E the fitting procedure and the values of the optimal parameters. Interestingly, we find a protein degradation time *τ*_*p*_ = 1h9min, corresponding to an average protein half-life time: *τ*_1*/*2_ ∼48min which is very close to the value 43min, measured in [33]. Moreover, we obtain a saturation time *t*^**^ = 2*h*44min which remarkably corresponds to the time at which the dry mass density starts to be diluted Fig.2.E, thus confirming the most critical prediction of our model. We can then test our predictions on the two other independent datasets at our disposal, i.e., the dry mass density, obtained from suspended microchannel resonator (SMR) experiments, and the normalized protein number, from fluorescent intensity measurements. We emphasize that the subsequent comparisons with experiments are done without any adjustable parameters. The agreement between theory and experiment is striking Fig.2.D,E, and gives credit to our model. We underline that the value of the density of water that we used is 4 % higher than the expected value, *ρ*_*w*_ = 1.04 kg*/*L to plot Fig.2.E. This slight difference originates from the fact that our simplified theory assumes that the dry mass is entirely due to proteins whereas proteins represent only 60% of the dry mass. This hypothesis is equivalent to renormalizing the density of water as shown in Appendix IV D.

In summary, our theoretical framework combining the PLM with a growth model and a model of amino-acid biosynthesis provides a consistent quantitative description of the dry mass density homeostasis and its subsequent dilution at senescence without invoking any genetic response of the cell; the dilution is due to the physical crowding of mRNAs by ribosomes. It also solves a seemingly apparent paradox stating that the volume is proportional to the number of proteins although their concentrations are low in the cell without invoking any non-linear term in the osmotic pressure (see Discussion and Appendix III L).

### F. Mitotic swelling

Our previous results explain well the origin of the dilution of the cellular dry mass at senescence. But can the same framework be used to understand the systematic dry mass dilution experienced by mammalian cells at mitotic entry ? Although this so called mitotic swelling or mitotic overshoot is believed to play a key role in the change of the physio-chemical properties of mitotic cells, its origin remains unclear [34],[35].

We first highlight five defining features of the mitotic overshoot. (1) It originates from an influx of water happening between prophase and metaphase, resulting in a typical 10% volume increase of the cells. (2) The swelling is reversible and cells shrink back to their initial volume between anaphase and telophase. (3) This phenomenon appears to be universal to mammalian cells, larger cells displaying larger swellings. (4) Cortical actin was shown not to be involved in the process, discarding a possible involvement of the mechanosensitivity of ion channels, contrary to the density increase observed during cell spreading [16] (5) Nuclear envelope breakdown (NEB) alone cannot explain the mitotic overshoot since most of the swelling is observed before the prometaphase where NEB occurs [34],[35].

The dry mass dilution at mitotic overshoot is thus different from the cases studied in the previous section. First, it happens during mitosis when the dry mass is constant [35]. Second, the 10% volume increase implies that we need to improve the simplified model used above, which considers only metabolites and proteins (and their counterions). Having in mind that ions play a key role in the determination of the cell volume Fig.1C, we show how every feature of the mitotic overshoot can be qualitatively explained by our theory, based on a well-known electrostatic property of charged polymer called counterion condensation first studied by Manning [36]. Many counterions are strongly bound to charged polymers (such as chromatin) because the electrostatic potential at their surface creates an attractive energy for the counterions much larger than the thermal energy *k*_*B*_*T*. The condensed counterions partially neutralize the charge of the object and reduce the electrostatic potential. Condensation occurs up to the point where the attractive energy for the free counterions is of the order *k*_*B*_*T*. The condensed counterions then do not contribute to the osmotic pressure given by Eq.2 which determines the cell volume. These condensed counterions act as an effective “internal” reservoir of osmolytes. A release of condensed counterions increases the number of free cellular osmolytes and thus the osmotic pressure inside the cell. Therefore, it would lead to an influx of water in order to restore osmotic balance at the plasma membrane Fig.3.

**FIG. 3.**
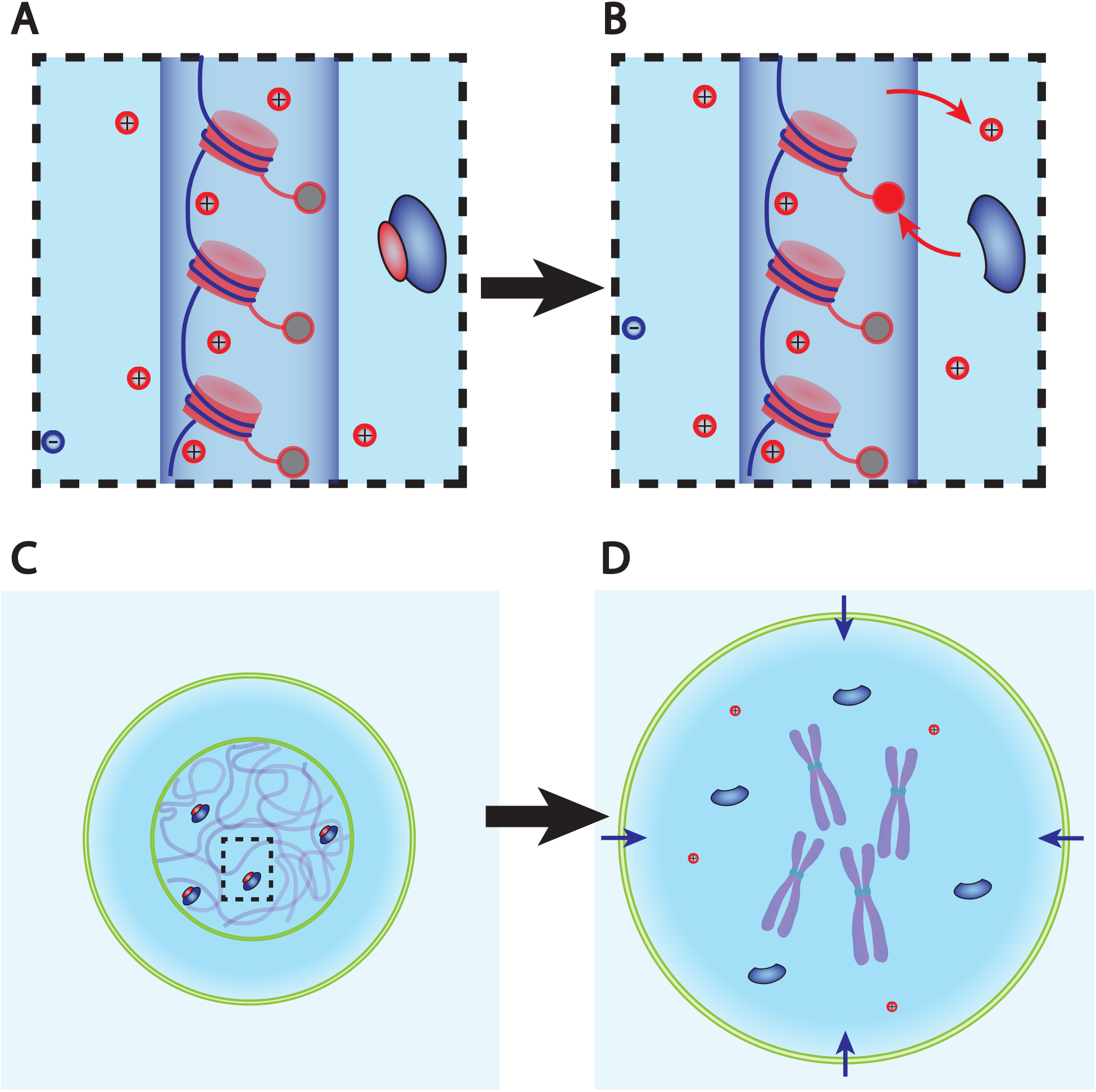
Dry mass dilution at mitosis is explained with the PLM by the decondensation of chromatin counterions following histone tail modifications. (A) and (B) Microscopic working model. An enzyme gives its positive charge to a histone, resulting in the release of a condensed counterions. Ions depicted within the chromatin (dark blue cylinder) are condensed and those outside are freely diffusing and participate in the nuclear osmotic pressure. (C) and (D) The subsequent increase in the number of osmolytes lead to a water influx in order to sustain osmotic balance at the plasma membrane of mammalian cells. For readability, other osmolytes are not displayed.

But how to explain such a counterion release at mitotic overshoot? For linear polymers such as DNA, the condensation only depends on a single Manning parameter 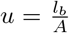; where, *l*_*b*_ is the Bjerrum length Tab.S1 which measures the strength of the coulombic interaction and A the average distance between two charges along the polymer. The crucial feature of Manning condensation is the increase of the distance between charges *A* by condensing counterions and thus effectively decreasing u down to its critical value equal to 1 (see Appendix V for a more precise derivation). Hence, the number of elementary charges carried by a polymer of length *L*_*tot*_ is 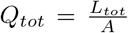 before condensation. After condensation, the effective distance between charges increases to *A*^*eff*^ = *A* · *u* such that the effective number of charges on the polymer is reduced to 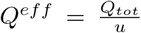. The number of counterions condensed on the polymer is 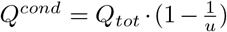. The most important consequence of these equations is that they suggest that a structural modification of the chromatin could lead to a counterion release. Indeed, making the chromatin less negatively charged, i.e., increasing A, is predicted to decrease u and thus to lead to the decrease of *Q*^*cond*^. Detailed numerical simulations of chromatin electrostatics show that this description is qualitatively correct [37].

Biologists have shown that chromatin undergoes large conformational changes at mitotic entry. One of them attracted our attention in light of the mechanism that we propose. It is widely accepted that the affinity between DNA and histones is enhanced during chromatin compaction by stronger electrostatic interactions thanks to specific covalent modifications of histone tails by enzymes. Some of these modifications such as the deacetylation of lysines add a positive charge to the histone tails, hence making the chromatin less negatively charged [26]. Moreover, histone tails are massively deacetylated during chromatin compaction [38], potentially meaning that this specific reaction plays an important role in counterion release and thus on the observed mitotic swelling. However, we underline that the idea that we propose is much more general and that any reaction modifying chromatin electrostatics is expected to impact the swelling. The question whether deacetylation of lysines is the dominant effect is left open here.

Is the proposed mechanism sufficient to explain the observed 10% volume increase? We estimate the effective charge of chromatin for a diploid mammalian cell to be *Q*^*eff*^ = 2 · 10^9^ e^−^ and the number of condensed monovalent counterions to be *Q*^*cond*^ = 8 · 10^9^ (see Appendix III). The PLM framework predicts the subsequent volume increase induced by the hypothetical release of all the condensed counterions of the chromatin. We find an increase of order Δ*V* ∼ 100 − 150*µ*m^3^ which typically represents 10% of a mammalian cell size (see Appendix III and Eq.S.26). Admittedly crude, this estimate suggest that chromatin counterion release can indeed explain the amplitude of mitotic swelling.

In summary, the combination of the PLM framework with a well-known polymer physics phenomenon allows us to closely recapitulate the features displayed during mitotic swelling. In brief, the decondensation of the chromatin condensed counterions, hypothetically due to histone tail modifications, is sufficient to induce a 10% swelling. This implies that, all mammalian cells swell during prophase and shrink during chromatin decondensation after anaphase; again, consistent with the dynamics of the mitotic overshoot observed on many cell types. Another salient implication is that the amplitude of the swelling is positively correlated with the genome content of the cells: cells having more chromatin are also expected to possess a larger “internal reservoir” of osmolytes, which can participate in decondensation. This provides a natural explanation for the observed larger swelling of larger cells. For instance, Hela cells were shown to swell on average by 20%, in agreement with the fact that many of them are tetraploid. Admittedly, many other parameters enter into account and may disrupt this correlation such as the degree of histone tail modifications or the initial state of chromatin; The existence of a larger osmolyte reservoir does not necessarily mean that more ions are released.

Finally, we point out that the ideas detailed in this section can be tested experimentally using existing in vivo or in vitro methods. For example, we propose to massively deacetylate lysines during interphase, by either inhibiting lysine acetyltransferases (KATs) or overexpressing lysine deacetylases (HDACS), in order to simulate the mitotic swelling outside mitosis. We also suggest to induce mitotic slippage or cytokinesis failure for several cell cycles, to increase the genome content, while recording the amplitude of swelling at each entry in mitosis [39].

### G. Nuclear scaling

Another widely documented scaling law related to cell volume states that the volume of cell organelles is proportional to cell volume ([40],[3]). As an example, we discuss here the nuclear volume. We develop a generalised “nested” PLM that explicitly accounts for the nuclear and plasma membranes (see Fig.4.A). Instead of writing one set of equation (Eq.1,2,3) between the interior and the exterior of the cell, we write the same equations both inside the cytoplasm and inside the nucleus (see Eq.S.52). Before solving this nonlinear system of equations using combined numerical and analytical approaches, we draw general conclusions imposed by their structure. As a thought experiment, we first discuss the regime where the nuclear envelope is not under tension so that the pressure jump at the nuclear envelope Δ*P*_*n*_ is much smaller than the osmotic pressure inside the cell Δ*P*_*n*_ *<<* Π_0_. The osmotic balance in each compartment implies that the two volumes have the same functional form as in the PLM, with two contributions: an excluded volume due to dry mass and a wet volume equal to the total number of particles inside the compartment divided by the external ion concentration (see Eq.S.53). It is noteworthy that the total cell volume, the sum of the nuclear and cytoplasmic volumes, is still given by Eq.4 as derived in the simple PLM. This result highlights the fact that the PLM strictly applies in the specific condition where the nuclear envelope is under weak tension. In addition, a crucial consequence of the osmotic balance condition at the NE is that it leads to a linear scaling relation between the volumes of the two compartments:

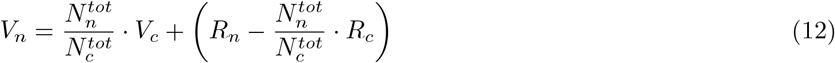

where *V*_*i*_, *R*_*i*_ and 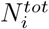 denote, respectively, the total volume, the dry volume and the total number of osmolytes of compartment i, the index *i* = *n, c* denoting either the nucleus *n*, or the cytoplasm *c*. Importantly, this linear scaling between the nucleus and the cytoplasm was reported repeatedly over the last century and is known as nuclear scaling [4], [3]. While this conclusion is emphasized in some recent papers [32], [41], we point out that Eq.12 is only a partial explanation of the nuclear scaling. Indeed, we still need to understand what cellular and nuclear properties makes the ratio 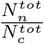 insensitive to external perturbations or to growth.

**FIG. 4.**
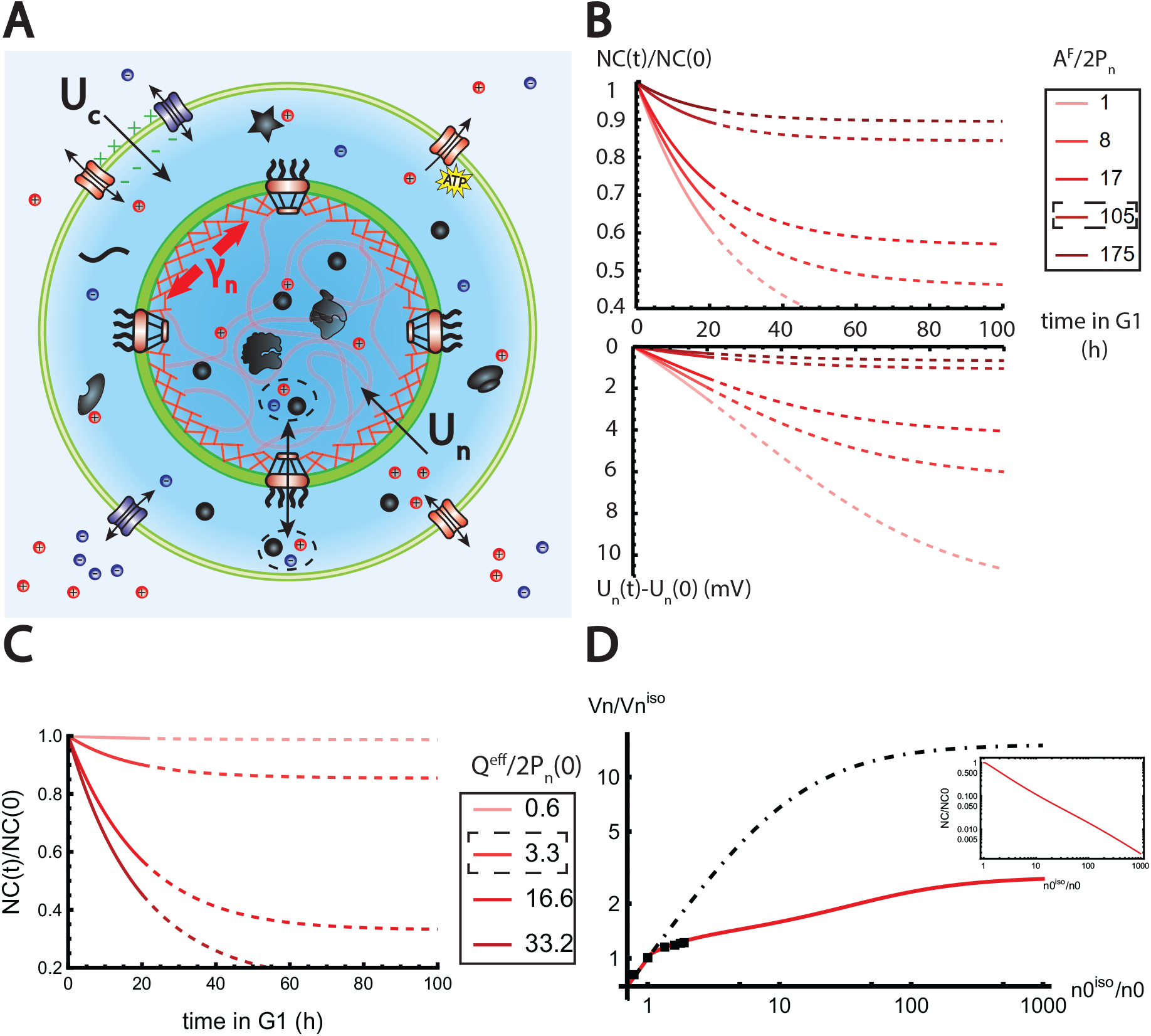
The nested PLM explains nuclear scaling. (A) Schematic of the nested PLM. Species in black are impermeant molecules (X) and are now partitioned between the cytoplasm and the nucleus. Among those, only metabolites (black circles) can cross the NE. The NE is composed of the membrane (green) and the lamina (red) can be stretched when the nuclear folds are flattened. (B) and (C) Simulations of the nested PLM Eq.S.52 during growth when the osmotic pressure is balanced at the NE. The growth rate was adjusted to data in [35] (B) Though permeable to the NE, Metabolites play a role in the homeostasis of the NC ratio by diluting chromatin (free) counterions which do not scale during growth (top plot). Higher variations of the NC ratio correlate with higher variations of the NE potential (bottom plot). (C) Variations of the NC ratio during growth for different chromatin charges. (D) Normalized nuclear volume after a hypo-osmotic shock. Nuclear volume saturates because of the tension at the NE, leading to the decrease of the NC ratio (inset: log-log plot). The dash-dotted line represents the nuclear volume if the number of osmolytes in the nucleus were assumed constant throughout the shock. Thus, showing that Metabolites leave the nucleus during the shock which strongly decreases nucleus swelling. The value at the saturations are given by Eq.17. The square black dots are data extracted from [42]. We used K = 50mN*/*m ^a^ and *s* = 4% folds to fit the data.

### H. NC ratio in the low tension regime

We now examine the influence of the various cell osmolytes on the NC ratio. For the sake of readability, we assume that the volume fraction occupied by the dry mass is the same in the nucleus and the cytoplasm (see Appendix VI A). The NC ratio is then the ratio between the wet volumes. Following the lines of our previous discussion, four different components play a role in volume regulation: chromatin (indirectly through its non-condensed counterions), proteins (mainly contributing to the dry volume), metabolites and ions (mainly contributing to the wet volume). These components do not play symmetric roles in the determination of the NC ratio. This originates from the fact that metabolites are permeable to the nuclear membrane and that chromatin, considered here as a gel, does not contribute directly to the ideal gas osmotic pressure because its translational entropy is vanishingly small [45]. The nested PLM leads to highly nonlinear equations that cannot be solved analytically in the general case (see Eq.S.52). Nevertheless, in the particular regime of monovalent osmolytes and high pumping *z*_*a*_ = 1, *z*_*p*_ = 1 and *α*_0_ = 0 corresponding to the case where there is no free anions in the cell, the equations simplify and are amenable to analytical results. This regime is physically relevant since it corresponds to values of the parameters close to the ones that we estimated (Fig.1). For clarity, we first restrict our discussion to this particular limit. We will also discuss both qualitatively and numerically the influence of a change of the parameters later. In this scenario, the nested PLM equations reduce to:

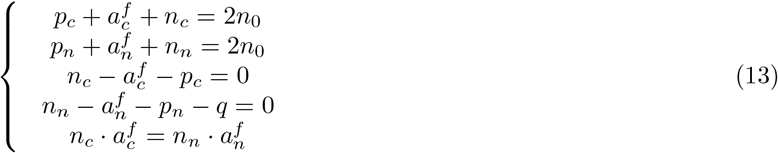

where the first and second equations correspond to osmotic pressure balance in the two compartments; the third and fourth equation correspond to macroscopic electroneutrality in each compartment; and the fifth equation is the balance of the chemical potential of the cations and metabolites on each side of the NE. *p*_*i*_, *n*_*i*_, *a*_*i*_ respectively accounts for the concentrations of proteins, cations and metabolites either in the cytoplasm - subscript c - or in the nucleus, subscript n. *q* accounts for the effective chromatin charge density. From these equations, we express the concentrations of cations in each compartment as functions of the extracellular concentration *n*_0_ and the chromatin charge density *q* (Eq.S.57), leading to the following expression of the NE potential:

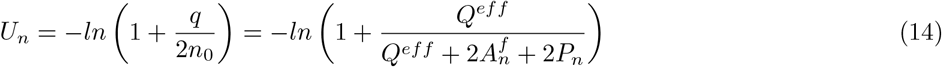

A salient observation from Eq.14 is that the NE potential difference *U*_*n*_ is a proxy of the chromatin charge density. At low *q, U*_*n*_ = 0, i.e., the respective concentrations of metabolites and cations are equal on each side of the membrane. Eq.13, also shows that the protein concentrations are equal in the two compartments. This implies that when the charge of chromatin is diluted, the volumes of the nucleus and of the cytoplasm adjust such that the NC ratio equals the ratio of protein numbers in the two compartments 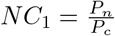. In the PLM, which considers a single compartment, a membrane potential appears as soon as there exist trapped particles in the compartment (see Appendix VI B and Eq.S.55). In contrast, our extended nested PLM predicts that in the case of two compartments, the system has enough degrees of freedom to adjust the volumes as long as *q* is small, thereby allowing the potential to be insensitive to the trapped charged proteins. At high values of the chromatin charge *Q*^*eff*^, *U*_*n*_ saturates to the value −*ln*(2) which in physical units is equivalent to −17mV at 300K. Note that this lower bound for the potential is sensitive to the average charge of the proteins *z*_*p*_ and can be lowered by decreasing this parameter. We also highlight that Eq.14 makes another testable prediction, namely, that the NE potential is independent of the external ion concentration. In the literature, NE potentials were recorded for several cell types [46]. They can vary substantially between cell types ranging from ∼ 0mV for Xenopus oocytes to −33mV for Hela Cells. This result is in line with our predictions. The Xenopus oocyte nucleus has a diameter roughly twenty times larger than typical somatic nuclei, but its chromatin content is similar [43], resulting in a very diluted chromatin and a vanishing NE potential. On the other hand, Hela cells are known to exhibit an abnormal polyploidy which may lead to a large chromatin charge density and a large nuclear membrane potential.

This last prediction allows to understand the influence of the metabolites on the NC ratio. An increase of the number of metabolites in the cell 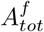, induces growth of the total volume (Eq.S.53), leading to the dilution of the chromatin charge and a strong decrease of the nuclear membrane potential (Eq.14). In the limit where 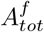 is dominant, we thus expect the NC ratio to be set to the value *NC*_1_. On the other hand, at low 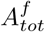, metabolites do not play any role on the NC ratio, which is then given by *NC*_2_ *> NC*_1_, with:

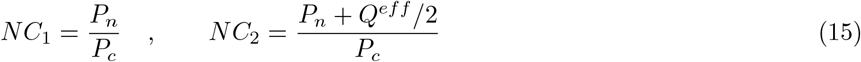

(see Eq.S.56 for the general formula). The actual NC ratio is intermediate between the two limiting behaviors (see Fig.S1B and Eq.S.62).

During cell growth, the ratio *NC*_1_ is constant, while the ratio *NC*_2_ varies with time. Indeed, if nucleo-cytoplasmic transport is faster than growth, the protein numbers *P*_*n*_ and *P*_*c*_ are both proportional to the number of ribosomes in the exponential growth regime and the ratio *NC*_1_ does not vary with time (see Appendix VI E). On the other hand, the DNA charge *Q*^*eff*^ is constant during G1 phase while *P*_*n*_ grows with time, so *NC*_2_ decreases with time. The fact that the NC ratio remains almost constant during growth ([47], [48]) suggest that cells are closer to the *NC*_1_ regime, and point at the crucial role of metabolites in setting the NC ratio (Fig.4 and S1.B). Importantly, these conclusions are overlooked in a large part of the existing literature ([31],[32],[41]) which often assumes that metabolites do not play any role on the NC ratio due to their permeability at the NE. We end this qualitative discussion by predicting the effect of a variation of the parameters *z*_*p*_, *z*_*a*_ and *α*_0_ that were so far assumed to be fixed. Our main point is that, any parameter change that tends to dilute the chromatin charge, also tends to increase the (negative) NE potential and make the NC ratio closer to the regime *NC*_1_ and further from the regime *NC*_2_. Consequently, increasing both *z*_*p*_ and *z*_*a*_, the number of counterions carried by each protein or metabolite increases, resulting in a global growth of the volume, hence to the dilution of the chromatin charge and to the increase of the NE potential. Any increase of the pumping parameter *α*_0_ (decrease of pumping efficiency) has a similar effect. It increases the number of ions in the cell resulting again in the dilution of the chromatin charge. Note that in the absence of pumping, (*α*_0_ = 1), the PLM predicts a diverging volume because this is the only way to enforce the balance of osmotic pressures at the plasma membrane (if there is no pressure difference at the membrane due to a cell wall).

Five crucial parameters have emerged from our analytical study: 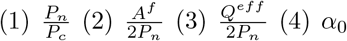 and (5) *z*_*p*_/*z*_*a*_. But what are the biological values of these parameters? We summarize our estimates in Appendix III. Importantly, the ratio between chromatin (free) counterions and the number of nuclear trapped proteins (and their counterions) is estimated to be of order one (see Appendix III and Fig.4.C). As a key consequence, we find that the NC ratio would be four times larger in the absence of metabolites Fig.S1.B. This non intuitive conclusion sheds light on the indirect, yet fundamental, role of metabolites on the NC ratio, which have been overlooked in the literature.

We now turn to a numerical solution to obtain the normalized variations of the NC ratio during growth in the G1 phase for different parameters Fig.4. Interestingly, variations of the NC ratio and variations of the NE potential are strongly correlated, a feature that can be tested experimentally Fig.4.B. Moreover, we deduce from our numerical results that, in order to maintain a constant NC ratio during the cell cycle, cells must contain a large pool of metabolites, see Fig.4.C. Our estimates point out that this regime is genuinely the biological regime, thus providing a natural explanation on the origin of the nuclear scaling, which is a robust feature throughout biology.

In summary, many of the predictions of our analysis can be tested experimentally. Experiments tailored to specifically modify the highlighted parameters are expected to change the NC ratio. For example, we predict that depleting the pool of metabolites, by modifying amino-acid biosynthesis pathways, i.e., lowering 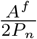, would lead to an increase of the NC ratio. Importantly, good metabolic targets in these experiments could be glutamate or glutamine because they account for a large proportion of the metabolites in the cell [28]. We also point out that cells with a smaller metabolic pool are expected to experience higher variations of the NC ratio during growth and thus larger fluctuations of this ratio at the population level Fig.4.B. These predictions could shed light on understanding the wide range of abnormal karyoplasmic ratio among cancer cells. Indeed, metabolic reprogramming is being recognized as a hallmark of cancer [49]; some cancer cells increase their consumption of the pool of glutamate and glutamine to fuel the TCA cycle and enhance their proliferation and invasiveness [50].

Moreover, disruption of either nuclear export or import is expected to change 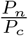 and thus the NC ratio. Numerical solutions of the equations displayed in Fig.S1 show a natural decrease of the NC ratio due to the disruption of nuclear import. On the other hand, if nuclear export is disrupted, we expect an increase of the NC ratio. This is in agreement with experiments done very recently in yeast cells [32]. The authors reported a transient decrease followed by an increase of the diffusivities in the nucleus. This is precisely what our theory predicts. The initial decay is due to the accumulation of proteins in the nucleus, resulting in an associated crowding. While, the following increase, is due to the impingement of ribosome synthesis. As this step requires nuclear export, it leads to the loss of the exponential growth and a decoupling between protein and amino acid numbers that drives the dilution of the nuclear content.

Finally, our framework also predicts that experiments that would maintain the 5 essential parameters unchanged, would preserve the nuclear scaling. We thus expect that, as long as the NE is not under strong tension, changing the external ion concentration does not influence the scaling directly. Experiments already published in the literature [15] shows precisely this feature.

## I. Mechanical role of the Lamina on the NC ratio

So far we have assumed that the osmotic pressure is balanced at the NE, which is a key condition for the linear relationship between nuclear and cytoplasmic volume. But why should this regime be so overly observed in biology? We first address this question qualitatively. For simplicity in the present discussion, we assume that DNA is diluted so that the NE potential is negligible. This implies that metabolites are partitioned so that their concentrations are equal in the nucleoplasm and the cytoplasm, hence cancelling their contribution to the osmotic pressure difference at the NE Eq.13. In the limit *α*_0_ ∼ 0, this allows to express the volume of the nucleus as:

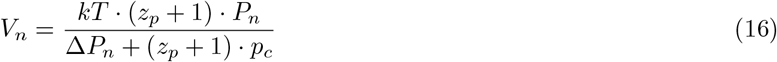

While the previous expression does not represent the exact solution of the equations, it qualitatively allows to realise that the NE hydrostatic pressure difference plays a role in the volume of the nucleus if it is comparable to the osmotic pressure exerted by proteins and their counterions. This pressure is in the 1000Pa range since protein concentration are estimated to be in the millimolar range (AppendixIII). We further estimated an upper bound for the nuclear pressure difference to be in the 10^4^Pa range (Eq.S.30). Admittedly crude, these estimates allow us to draw a threefold conclusion. (1) The nuclear pressure difference can be higher than the cytoplasmic pressure difference, in part due to the fact that Lamina has very different properties compared to cortical actin: it is much stiffer and its turnover rate is lower. This points out the possible role of nuclear mechanics in the determination of the nuclear volume contrary to the cortical actin of mammalian cells that does not play any direct role for the cell volume, (2) The typical hydrostatic pressure difference at which mechanical effects become relevant is at least two orders of magnitude lower for the nucleus than for the cytoplasm, for which it is of order *π*_0_, (3) Assuming linear elasticity, small NE extensions of 10% would be sufficient to impact nuclear volume. These conclusions stand in stark contrast to the observed robustness of the nuclear scaling, thus pointing out that the constitutive equation for the tension in the lamina is nonlinear. Biologically, we postulate that this non-linearity originates from the folds and wrinkles that many nuclei exhibit [22]. These folds could indeed play the effective role of membrane reservoirs, preventing the NE tension to grow with the nuclear volume, and setting the nuclear pressure difference to a small constant value, thereby maintaining cells in the scaling regime discussed in the previous sections. This conclusion is consistent with the results of Ref. [42], which observed that the nucleus exhibits non-linear osmotic properties.

To further confirm our conclusions quantitatively, we consider the thought experiment of non-adhered cells experiencing hypoosmotic shock. This experiment is well adapted to study the mechanical role of nuclear components on nuclear volume because it tends to dilute the protein content while increasing the hydrostatic pressure by putting the NE under tension. For simplicity, we ignore the mechanical contribution of chromatin that was shown to play a negligible role on nuclear mechanics for moderate extensions [51]. To gain insight into the non-linear set of equations, we split the problem into two parts. First, we identify analytically the different limiting regimes of nuclear volume upon variation of the number of impermeant molecules *X*_*n*_ present in the nucleus and the NE tension *γ*_*n*_. We summarize our results in a phase portrait (see Appendix VI F and Fig.S1). Two sets of regimes emerge: those, studied above, where nuclear and cytoplasmic osmotic pressures are balanced, and those where the nuclear hydrostatic pressure matters. In the latter situations, the nuclear volume does not depend on the external concentration and saturates to the value (see Appendix VI G) :

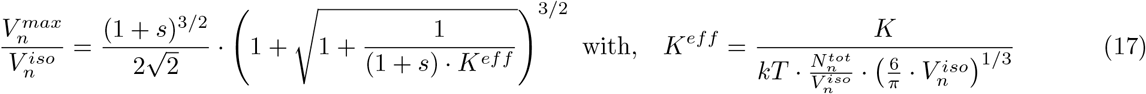

where, *s* and 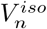 are respectively the fraction of membrane stored in the folds and the volume of the nucleus at the isotonic external osmolarity 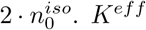 is an effective adimensional modulus comparing the stretching modulus of the nuclear envelope *K* with an osmotic tension that depends on the total number of free osmolytes contained by the nucleus 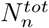. The saturation of the nuclear volume under strong hypoosmotic shock originating from the pressure build up in the nucleus after the unfolding of the folds, implies a significant decrease of the NC ratio and a loss of nuclear scaling Fig.4.D.

As a second step, we investigate the variations of 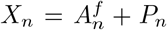 after the shock. Our numerical solution again highlight the primary importance of considering the metabolites 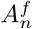 for the modelling of nuclear volume. Indeed, disregarding their contribution would lead to an overestimation of the number of trapped proteins. Additionally, *X*_*n*_ would remain constant during the osmotic shock, resulting in the reduction of the effective modulus of the envelope Eq.17. We would thereby overestimate the nuclear volume (Fig.4.D dashed line). In reality, since free osmolytes are mainly accounted for by metabolites which are permeable to the NE, the number of free osmolytes in the nucleus decreases strongly during the shock. This decrease can easily be captured in the limit where metabolites are uncharged *z*_*a*_ = 0. The balance of concentrations of metabolites Eq.13 implies that the number of free metabolites in the nucleus, 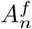, passively adjusts to the NC ratio:

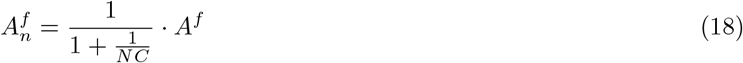

As mentioned earlier, the tension of the envelope is responsible for the decrease of the NC ratio. This in turn decreases the number of metabolites inside the nucleus, reinforcing the effect and thus leading to a smaller nuclear volume at saturation Fig.4.D. We find the analytical value of the real saturation by using Eq.17 with 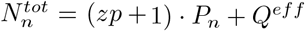, i.e., no metabolites remaining in the nucleus.

Our investigations on the influence of the hydrostatic pressure term in the nested PLM, lead us to identify another key condition to the nuclear scaling, i.e., the presence of folds at the NE. Moreover, although not the purpose of the present article, using our model to analyse hypoosmotic shock experiments could allow a precise charaterisation of the nucleus mechanics.

## III. DISCUSSION

In this study, we have investigated the emergence of the cell size scaling laws, which are the linear relations between dry mass, nuclear size and cell size, and which seem ubiquitous in living systems. Using a combination of physical arguments ranging from thermodynamics, statistical physics, polymer physics, mechanics and electrostatics, we have provided evidence that the robustness of these scaling laws arises from three physical properties : electroneutrality, balance of water chemical potential, and balance of ionic fluxes. The set of associated equations defines a model developed 60 years ago named the PLM. The major challenge in probing the origin of the scaling laws using the PLM, which we have addressed in this study, is to link a wide range of cell constituents and microscopic biological factors, such as ion transport, translation, transcription, chromatin condensation, nuclear mechanics, to the mesoscopic parameters of the PLM, Fig.1.B. A host of experimental papers has gathered evidence on these scaling laws and their breakdown over the past century [2],[3],[4], but no theoretical analyses have unified these observations within a single theoretical framework.

In order to go in this direction, we have simplified the PLM to its utmost based on the determination of precise orders of magnitude of the relevant parameters. The use of a simplified model focusing on the leading order effects, such as the homeostasis between amino-acids and proteins, is a powerful way to isolate and better study the origin of the scaling laws. This is embodied in the accurate predictions, without any adjustable parameters, for the dry mass dilution and the protein dynamics of yeast cells, which are prevented from dividing. A phenomenon that was so far unexplained [10] despite the fact that it is believed to be of fundamental biological importance [11] by establishing a functional relationship between cell size (and density) and cell senescence, potentially providing a novel mechanism driving this important aging process.

The key ingredient of our model is the consideration of small osmolytes and in particular metabolites and small ions. Their high number fractions among cell free osmolytes implies that they dominate the control of cell volume. We make three quantitative predictions from this finding (1) The homeostasis between amino-acids and proteins, originating from the enzymatic control of the amino-acid pool, explains the dry mass density homeostasis. The disruption of homeostasis, due to mRNA crowding by ribosomes or pharmacological treatment such as rapamycin, is predicted to lead to dry mass dilution upon cell growth, due to the saturation of the protein content while the number of amino-acids and thus the volume keeps increasing with time, (2) The dry mass dilution observed at mitotic entry for mammalian cells can naturally be explained by the release of counterions condensed on the chromatin, leading to the increase of the number of osmolytes inside the cell and to the subsequent influx of water to ensure osmotic pressure balance at the plasma membrane, (3) The robustness of the NC ratio to the predicted value 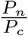 is due to the high pool of metabolites within cells, resulting in the dilution of the chromatin (free) counterions which do not scale during growth.

Interestingly, only few amino-acids represent most of the pool of the metabolites possessed by the cell, i.e., glutamate, glutamine and aspartate. Emphasizing their crucial role on cell size. Our investigations thus link two seemingly distinct hallmark of cancers : the disruption of the cell size scaling laws such as the abnormal karyoplasmic ratio, historically used to diagnose cancer, and metabolic reprogramming, some cancer cells showing an increased consumption of their pool of glutamate and glutamine to fuel the TCA cycle and enhance their proliferation and invasiveness [50]. This may thus represent possible avenues for future research related to the variability of nucleus size in cancer cells [52]. Moreover, the large pool of metabolites is a robust feature throughout biology [28], making it one of the main causes of the universality of the cell size scaling laws observed in yeasts, bacteria and mammalian cells. We believe that the more systematic consideration of such small osmolytes will allow to understand non-trivial observations. For instance, the recent observation of the increase of diffusivities in the nucleus after blocking nuclear export, is explained in our model by the decoupling between protein and amino-acid homeostasis after the impingement of ribosome synthesis, a step that requires nuclear export [32].

### A. The nucleoskeletal theory

To study the nuclear scaling law, we developed a model for nuclear volume, by generalizing the PLM, that includes both nuclear mechanics, electrostatics and four different classes of osmolytes. The clear distinction between these classes of components is crucial according to our analysis and is new. (1) Chromatin, considered as a gel, does not play a direct role in the osmotic pressure balance because its translational entropy is vanishingly small. Yet, it plays an indirect role on nuclear volume through its counterions. This creates an asymmetry in our system of equations, leading to the unbalance of ionic concentrations across the NE and to the appearance of a NE potential related to the density of chromatin. (2) Proteins, are considered as trapped in the nucleus, their number being actively regulated by nucleo-cytoplasmic transport. (3) Metabolites, are considered as freely diffusable osmolytes through the NE but not through the plasma membrane. Note that only half of the proteins are trapped in the nucleus because about half of them have a mass smaller than the critical value 30-60kDa [24], which corresponds to the typical cut-off at which they cannot freely cross nuclear pore complexes. This represents more a semantic issue than a physical one, and permeant proteins are rigorously taken into account as metabolites in the model, but are negligible in practice due to the larger pool of metabolites. (4) Free ions, are able to diffuse through the plasma membrane and the NE.

As a consequence, we show that the nuclear scaling originates from two features. The first one is the balance of osmotic pressures at the NE, that we interpret as the result of the non-linear elastic properties of the nucleus likely due to the presence of folds in the nuclear membrane of mammalian cells. Interestingly, yeast cells do not possess lamina such that the presence of nuclear folds may not be required for the scaling. In this regard, our model adds to a recently growing body of evidence suggesting that the osmotic pressure is balanced at the NE in isotonic conditions [41],[32],[42]. The second feature is the presence of the large pool of metabolites accounting for most of the volume of the nucleus. This explains why nuclear scaling happens during growth while the number of chromatin counterions does not grow with cell size.

Interestingly, although not the direct purpose of this article, our model offers a natural theoretical framework to shed light on the debated nucleoskeletal theory [4], [3]. Our results indicate that the genome size directly impacts the nuclear volume only if the number of (free) counterions of chromatin dominates the number of trapped proteins and the number of metabolites inside the nucleus. We estimate that this number is comparable to the number of trapped proteins while it is about 60 times smaller than the number of metabolites, in agreement with recent observations that genome content does not directly determine nuclear volume [3]. Although not directly, chromatin content still influences nuclear volume. Indeed, nuclear volume (Eq.S.58) is mainly accounted by the number of metabolites, which passively adjusts according to Eq.S.52(7). In the simple case, of diluted chromatin and no NE potential, metabolite concentration is balanced and 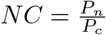, such that the metabolite number depends on two factors (Eq.18). The first one, is the partitioning of proteins 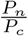, that is biologically ruled by nucleo-cytoplasmic transport in agreement with experiments that suggest that the nucleo-cytoplasmic transport is essential to the homeostasis of the NC ratio [3]. The second one is the total number of metabolites, ruled by the metabolism Eq.9, which ultimately depends upon gene expression (Appendix IV), as shown by genetic screen experiments done on fission yeast mutants [3]. However, when the chromatin charge is not diluted, which is likely to occur for cells exhibiting high NE potential such as some cancer cells, our theory predicts that the number of metabolites in the nucleus also directly depends on the chromatin content due to electrostatic effects. This highlights the likely importance of chromatin charge in the nuclear scaling breakdown in cancer.

### B. Role of NE breakdown in cell volume variations

The nested PLM predicts that the cell swells upon NE breakdown if the NE is under tension. NE breakdown occurs at prometaphase, and does not explain most of the mitotic swelling observed in [34, 35], which occurs at prophase. Within our model based on counterion release, mitotic swelling is either associated with cytoplasm swelling if the released counterions leave the nucleus, or with nuclear swelling if they remain inside. In the latter case, swelling at prophase would be hindered by an increase of NE tension, and additional swelling would occur at NE breakdown. This prediction can be tested by artificially increasing the NE tension through strong uniaxal cell confinement [53], which would synchronise mitotic swelling with NE breakdown.

### C. Physical grounds of the model

Physically, why can such a wide range of biological phenomena be explained such a simple theory? A first approximation is that we calculated the osmotic pressure considering that both the cytoplasm and the nucleus are ideal solutions. However, it is known that the cytoplasm and the nucleoplasm are crowded ([54],[55]). The qualitative answer again comes from the fact that small osmolytes constitute the major part of the free osmolytes in a cell so that steric and short range attractive interactions are only a small correction to the osmotic pressure. We confirm this point by estimating the second virial coefficient that gives a contribution to the osmotic pressure only of order 2kPa (see Appendix III), typically 2 orders of magnitude smaller than the ideal solution terms Fig.1.B. However, note that we still effectively take into account excluded volume interactions in our theory through the dry volume *R*. Moreover, we show in appendix VII) that although we use an ideal gas law for the osmotic pressure, the Donnan equilibrium effectively accounts for the electrostatic interactions. Finally, our theory can be generalized to take into account any ions species and ion transport law while keeping the same functional form for the expressions of the volume Eq.S.18, as long as only monovalent ions are considered. This is a very robust approximation, because multivalent ions such as calcium are in the micromolar range. Together, these observations confirm that the minimal formulation of the PLM that we purposely designed is well adapted to study cell size.

### D. Future extensions of the theory

As a logical extension of our results, we suggest that our framework be used to explain the scaling of other membrane bound organelles such as vacuoles and mitochondria [40]. We show in appendix (Eq.S.80) that the incorporation of other organelles into our framework lead to the same equations as for the nucleus, thus pointing out that the origin of the scaling of other organelles may also arise from the balance of osmotic pressures. We also propose that our theory be used to explain the scaling of membraneless organelles such as nucleoids [56]. Indeed, the Donnan picture that we are using does not require membranes [57]. However, we would have to add other physical effects in order to explain the partitioning of proteins between the nucleoid and the bacterioplasm.

Taken as a whole, our study demonstrates that cell size scaling laws can be understood and predicted quantitatively on the basis of a remarkably simple set of physical laws ruling cell size as well as a simple set of universal biological features. The multiple unexplained biological phenomena that our approach allows to understand indicates that this theoretical framework is fundamental to cell biology and will likely benefit the large community of biologists working on cell size and growth.

## IV. ACKOWLEDGMENTS

We thank Matthieu Piel and the members of his team, in particular Damien Cuvelier, Alice Williart, Guilherme Nader and Nishit Srivastava, for insightful discussions and for showing us data that originally motivated the theory on nuclear volume; Thomas Lecuit for introducing us to the cell size scaling laws with his 2020 course at College de France entitled “Volume cellulaire determinants Physico-chimiques et regulation”; Pierre Recho for showing us his seminal work on nuclear volume; the members of the UMR 168, Amit Singh Vishen, Ander Movilla, Sam Bell, Mathieu Dedenon and Joanna Podkalicka as well as Dan Deviri for fruitful discussions. The Sens laboratory is a member of the Cell(n)Scale Labex.

## Footnotes

a The value of K used to fit the data [42] is twice the measured value in [43]. The rationale is threefold. (1) Nuclei used in [42] are chondrocite nuclei originating from articular cartilage. They possess a high Lamina A to Lamina B ratio and are thus likely to be stiffer [44] (2) We could lower the value of the fitted K by increasing the pumping efficiency *α*_0_. A more detailed caracterisation of the PLM parameters for chondrocites would be required to precisely infer the elastic properties of the NE. (3) Considering the chromatin mechanical contribution would increase K by a factor *E*_*DNA*_ *· R*_*nucleus*_; with *E*_*DNA*_ the elastic modulus of the chromatin and *R*_*nucleus*_ the radius of the nucleus.

