## Supplementary for "Cell size scaling laws: a unified theory"


### I. PLM FUNDAMENTAL EQUATIONS

In this section we derive and discuss the three physical constraints we used throughout our paper to study cell volume regulation. These results are classical and can be found in a reference textbook such as [1].

#### A. Electroneutrality

The intrinsic length scale associated to the Poisson equation is the Debye length. It appears explicitly in the linearized version of the Poisson equation also called the Debye-Huckel equation. It reads:

$$\lambda_D = \left(\frac{1}{4\pi \cdot l_b \cdot (n^+ + n^-)}\right)^{\frac{1}{2}} \tag{S.1}$$

Where: $l_B = \frac{e^2}{4\pi k T \epsilon_r \epsilon_0}$ is the Bjerrum length - which qualitatively corresponds to the distance between two elementary charges at which the electrostatic energy will be comparable to the thermal energy. $l_B \approx 0.7$nm in water at 300K. In the unit used in this paper (concentrations in mMol) the Debye length can be estimated using the following formula:

$$\lambda_D \approx \frac{9.7}{\sqrt{n^+(mM) + n^-(mM)}} \cdot nm \tag{S.2}$$

For a typical mammalian cell $n^+(\text{mM}) + n^-(\text{mM}) \approx 180\text{mM}$ (Fig.1) which leads to a Debye length $\lambda_D \approx 0.7$nm. Thus, the Debye length is at least 3 orders of magnitude smaller than the typical radius of a cell or of a nucleus. This justifies the approximation of electroneutrality used throughout the main paper for length scales much larger than the Debye length.

#### B. Balance of water chemical potential

We define the osmotic pressure as:

$$\Pi = -\frac{1}{v_w} \cdot (\mu_w - \mu_w^*) \tag{S.3}$$

where : $v_w$ is the molecular volume of water, $\mu_w$, $\mu_w^*$ are the chemical potential of water respectively in the real solution and in a pure water solution. Assuming that water is incompressible, $\mu_w^* = \mu_0(T) + v_w \cdot P$, where $P$ is the hydrostatic pressure. When water is equilibrated, thermodynamics imposes that the chemical potential of water is equal on both sides of the membrane. From the previous equations it is straightforward to derive Eq.2 in the main text.


*
†
‡

#### C. Balance of ionic fluxes

The total flux J of cations - respectively anions - is decomposed between 3 main contributions: active pumping, electrical conduction and entropic diffusion. For simplicity we assumed that only cations are pumped out of the cell. This simplifying choice was made to model the Na/K pump which is one of the most relevant cationic pumps. Though, we show in subsection.II that this assumption is not critical since the equations keep the same functional form if it is relaxed. As a convention, we choose $J$ to be positive when ions are entering the cell. At steady-state, the fluxes vanish:

$$\begin{cases} J_{+,tot} = g^+ \cdot \left[ -e \cdot U_c - kT \cdot ln\left(\frac{n^+}{n_0}\right)\right] - p = 0 \\ J_{-,tot} = g^- \cdot \left[ e \cdot U_c - kT \cdot ln\left(\frac{n^-}{n_0}\right)\right] = 0 \end{cases} \tag{S.4}$$

where $p$ is the pumping flux, $g^{\pm}$ are the membrane conductivities for cations and anions, and $U_c$ is the cell transmembrane potential, which can be expressed as:

$$U_c = -\frac{kT}{e} \cdot ln\left(\frac{n^+}{n_0}\right) - \frac{p}{g_+} = \frac{kT}{e} \cdot ln\left(\frac{n^-}{n_0}\right) \tag{S.5}$$

For Eq.S.5 to be verified, the following relationship between $n^+$ and $n^-$ must be imposed :

$$\begin{cases} n^+ \cdot n^- = \alpha_0 \cdot n_0^2 \\ \alpha_0 = e^{-\frac{p}{g^+}} \end{cases} \tag{S.6}$$

The latter equation takes the form of a generalized Donnan ratio that includes the active pumping of cations. The usual Donan ratio [1] is recovered when $p = 0$. The generalized Donnan ratio Eq.3 together with the electroneutrality condition Eq.1 yield analytic expressions for the ionic densities $n^+$ and $n^-$ (the notations are defined in the main text):

$$\begin{cases} n^+ = \frac{zx + \sqrt{(zx)^2 + 4\alpha_0 n_0^2}}{2} \\ n^- = \frac{-zx + \sqrt{(zx)^2 + 4\alpha_0 n_0^2}}{2} \end{cases} \tag{S.7}$$

The cell osmotic pressure can thus be expressed as:

$$\frac{\pi}{kT} = \sqrt{(zx)^2 + 4\alpha_0 n_0^2} + x \tag{S.8}$$

### II. GENERAL EXPRESSIONS OF THE VOLUME IN THE PLM MODEL

The system of equation formed by Eq.1,2,3 is nonlinear and cannot be solved analytically in its full generality. One complexity arises from the difference of hydrostatic pressure $\Delta P$. Intuitively, if the volume increases, the surface increases which may in some situations increase the tension of the envelope and in turn impede the volume growth. Mathematically, Laplace law relates the difference of hydrostatic pressure to the tension $\gamma$ and the mean curvature of the interface - which simplifies to the radius of the cell R in a spherical geometry. The difference of hydrostatic pressure then reads:

$$\Delta P = \frac{2\gamma}{R} \tag{S.9}$$

In the case where the interface exhibits a constitutive law which is elastic $\gamma = K \cdot \frac{S-S_0}{S_0}$, it is easy to see that $\Delta P$ exhibits power of V which makes the problem non analytical. However, we can get around this limitation in two biologically relevant situations :

- When $\Delta P$ is negligible. As shown in Fig1.B this happens for mammalian cells that do not possess cellular walls.
- When $\Delta P$ is buffered by biological processes. We argue that this situation applies for yeasts and bacteria during growth. Indeed, if the volume increase is sufficiently slow, one can hypothesize that cells have time to add materials to their cellular walls such that the tension does not increase during growth.

We give the corresponding analytical expressions under these two hypotheses in the next two paragraphs.

#### A. Analytical expression of the volume when hydrostatic pressure difference is negligible

The balance of water chemical potential Eq.2, neglecting the difference of pressure and injecting the expressions for the ionic densities Eq.S.7 leads to the following equation for the density of impermeant molecules $x$ :

$$(z^2-1)\cdot x^2 + 4n_0\cdot x - 4\cdot(1-\alpha_0)\cdot n_0^2 = 0 \tag{S.10}$$

Solving this equation and using the definition of the density of impermeant molecules $x = \frac{X}{V-R}$ yield the expression for the volume of the cytoplasm :

$$\begin{cases} V - R = \frac{kT\cdot N^{tot}}{\Pi_0} \\ N^{tot} = X\cdot\frac{(z^2-1)}{-1+\sqrt{1+(1-\alpha_0)(z^2-1)}} \end{cases} \tag{S.11}$$

The volume can thus be written as an ideal gas law with a totel number of free osmolytes $N^{tot}$. This number takes into account the different ions and is thus larger than the actual number of impermeant molecules $X$. In the limit of very fast pumping - $\alpha_0 \to 0$ - Eq.S.11 reduces to the expression given in the main text Eq.4

#### B. Analytical expression of the volume when $\Delta P$ is buffered

The same procedure can be used when $\Delta P$ is buffered (independent of the volume). The final expression reads:

$$\begin{cases} V - R = \frac{kT\cdot N^{tot}(\Delta P)}{(\pi_0+\Delta P)} \\ N^{tot}(\Delta P) = X\cdot\frac{z^2-1}{-1+\sqrt{1+(z^2-1)\cdot\left(1-\frac{\alpha_0}{\left(1+\frac{\Delta P}{kT\cdot 2n_0}\right)^2}\right)}} \end{cases} \tag{S.12}$$

Interestingly, the wet volume $V-R$ remains proportional to the number of impermeant molecules $X$ in this limit.

#### C. Analytical expression of the volume for an arbitrary number of ions and active transports

In this subsection, we generalize the PLM to any type of ions and any ionic transport. Each ion can be actively transported throughout the membrane. Importantly, we show that - as long as ions are monovalent - the PLM equations and solutions take the same functional form as the two-ions model used in the main text. We use the same notations as in the main text (Section II A and Fig.1), except that we now add subscript $i$ to refer to the ion of type $i$. For instance, $z_i^-$ - respectively $n_i^-$ - refers to the valancy - respectively the concentration - of the anion $i$. The densities of positive / negative charges in the cell read:

$$\begin{cases} d^+ = \sum_j z_j^+\cdot n_j^+ \\ d^- = \sum_j z_j^-\cdot n_j^- \end{cases} \tag{S.13}$$

Electroneutrality thus simply reads:

$$d^+ - d^- - z \cdot x = 0 \tag{S.14}$$

Balancing ionic fluxes for each ion types, as in Eq.S.4, leads to :

$$\begin{cases} n_j^{+/-} = n_j^0 \cdot \alpha_j \cdot e^{(-/+)\cdot z_j \cdot \frac{e \cdot U_c}{kT}} \\ \alpha_j = e^{-\frac{p_j}{g_j}} \end{cases} \tag{S.15}$$

Using Eqs.S.13,S.14, S.15 and assuming that all ions are monovalent, the product of the cationic and anionic densities can be expressed as :

$$d^+ \cdot d^- = \underbrace{\left(\sum_j n_j^{+,0} \cdot \alpha_j\right) \cdot \left(\sum_i n_i^{-,0} \cdot \alpha_i\right)}_{\underset{def}{\equiv} \tilde{\alpha}(n_i^0)} \tag{S.16}$$

and the analytical solution of the full problem reads:

$$\begin{cases} d^+ = \frac{zx + \sqrt{(zx)^2 + 4\tilde{\alpha}(n_i^0)}}{2} \\ \\ d^- = \frac{-zx + \sqrt{(zx)^2 + 4\tilde{\alpha}(n_i^0)}}{2} \end{cases} \tag{S.17}$$

$$\begin{cases} V - R = \frac{kT \cdot N^{tot}(\Delta P)}{(\pi_0 + \Delta P)} \\ \\ N^{tot}(\Delta P) = X \cdot \frac{z^2 - 1}{-1 + \sqrt{1 + (z^2 - 1) \cdot \left(1 - \frac{4\tilde{\alpha}(n_i^0)}{\left(\frac{1}{kT} \cdot \Pi_0 + \frac{1}{kT} \cdot \Delta P\right)^2}\right)}} \end{cases} \tag{S.18}$$

which shows a similar form as the two-ion model, Eqs.S.7,S.12.

### III. ORDER OF MAGNITUDES

Throughout the main text, we used order of magnitudes to guide our investigations and justify our approximations. For the sake of readability we gather all the parameter significations, values, and origins in Table S1.

#### A. Protein concentration

We use data published in [2] to estimate the typical concentration of proteins in mammalian cell $p_{tot}$ as :

$$p_{tot} = \%_p^{mass} \cdot \frac{\rho}{\mathcal{M}_a \cdot l_p \cdot (1 - \frac{R}{V})} \sim 2mMol \tag{S.19}$$

where, $\%_p^{mass}$ is the fraction of dry mass occupied by proteins Fig.1.D.

#### B. mRNA to protein fraction

In figure 1.C we neglected the contribution of mRNAs to the wet volume of the cell. The rationale behind this choice is twofold. (1) Proteins represent less than 1% of the wet volume (2) The mRNA to protein number fraction is estimated to be small, due to the fact that the mass of one mRNA is 9 times greater than the one of a protein while the measured fraction of mRNA to dry mass is of the order 1% [3] :

$$\frac{M_{tot}}{P_{tot}} = \frac{\mathcal{M}_p}{\mathcal{M}_{mRNA}} \cdot \frac{\%^{mass}_{mRNA}}{\%^{mass}_p} \sim \frac{1}{500} \tag{S.20}$$

Thus, mRNAs contribute even less than proteins to the wet volume.

#### C. Metabolite concentration

We find the metabolite concentration self-consistently by enforcing balance of osmotic pressure at the plasma membrane Eq.2 :

$$a^f = 2n_0 - p_{tot} - n^+ - n^- \sim 118 mMol \tag{S.21}$$

where the concentrations of ions were reported in [3] (see Fig.1). This high value of metabolite concentration is coherent with reported measurements [4].

#### D. Contribution of osmolytes to the wet volume of the cell Fig.1.C

The contribution of osmolytes to the wet volume fraction is simply equal to the ratio of the osmolyte concentration to the external osmotic pressure, here equal to $2n_0$ Eq.4. The concentration of specific amino-acids and metabolites were estimated using their measured proportion in the metabolite pool [4] times the total concentration of metabolites $a^f$ (Eq.S.21).

#### E. Amino-acids contribution to the dry mass

One of the main conclusions from our order of magnitude estimates is that amino-acids play an essential role in controlling the volume but have a negligible contribution to the cell's dry mass. This originates from the large average size of proteins $l_p \sim 400$a.a. The contribution of amino-acids to the dry mass reads:

$$\%^{mass}_{a.a} = \%^{number}_{a.a} \cdot \frac{a^f}{p_{tot}} \cdot \frac{\%^{mass}_p}{l_p} \sim 6\% \tag{S.22}$$

where, $\%^{number}_{a.a} \sim 73\%$ is the number fraction of amino-acids among metabolites.

#### F. Effective charge of chromatin

The average effective charge per nucleosome is estimated to be:

$$Q^{eff}_{pernucleosome} = L_{Link} \cdot \frac{Q_{bp}}{u_{DNA}} + L_{wrap} \cdot Q_{bp} - Q_{hist} - Q_{wrap} = 71 \tag{S.23}$$

where, the right hand side can be understood as the total negative charge of pure DNA, screened in part by histone positive charges and by the manning condensed counterions. Note that the number of condensed counterions $Q_{wrap}$ around the wrapped DNA simulated in [5] is similar to the value expected by the manning theory which we estimate to be 164 elementary charges.

The number of nucleosomes is simply $N_{hist} = \frac{L_{tot}}{L_{nucleosomes}} = 3 \cdot 10^7$ such that the effective charge of chromatin is estimated to be :

$$Q^{eff} = 2 \cdot 10^9 \tag{S.24}$$

#### G. Condensed counterions on chromatin

The condensed counterions on chromatin can simply be found from the effective charge of the chromatin, the total charge of pure DNA and the charge of histones. We obtain:

$$Q^{cond} = Q_{bp} \cdot L_{tot} - Q^{eff} - Q_{hist} \cdot N_{hist} \sim 8 \cdot 10^9 \tag{S.25}$$

#### H. Estimation of the amplitude of the Mitotic Swelling

At mitosis cells have doubled their genome content such that we double the number of condensed counterions estimated earlier for a diploid mammalian cell. Using the PLM, we compute the amplitude of swelling if all the chromatin condensed counterions were released at the same time, assuming an external osmolarity of $n_0 = 100 - 150$mM.

$$\Delta V = \frac{2 \cdot Q^{cond}}{2n_0} \sim 100\mu\text{m}^3 \tag{S.26}$$

Note that $\Delta V$ must scale with the number of genome duplications. For instance, for tetraploid cells, the previous amplitude must be doubled.

#### I. Average charge of proteins and metabolites

The average charge of proteins used in the paper, $z_p \sim 0.8$, was estimated from [6] assuming that Histidines are neutral. This is reasonable because their Pka is of order 6 while typical physiological pH is of order 7.4, so that only 4% of histidines are charged in the cell.

The average charge of metabolites is assumed to be $z_a \sim 1$ since glutamate is the most abundant [4]. We have checked that changing this parameter does not alter our conclusions.

#### J. Absolute number of osmolytes

To obtain the Figure.4, we had to estimate the parameter $NC_1 = \frac{P_n}{P_c}$ and thus, the number of protein trapped inside the nucleus $P_n$ at the beginning of interphase. The total number of proteins and metabolites at the beginning of interphase is simply obtained by multiplying the concentration of proteins $p_{tot}$, $m_{tot}$ estimated earlier by the volume at the beginning of interphase, measured to be equal to 1250$\mu$m$^3$, minus the dry volume which roughly represent 30% of the total volume [2] and Fig.1.B.

$$P_{tot} = p_{tot} \cdot (V - R) \sim 10^9 \tag{S.27}$$

$$A^f = \frac{a^f}{p_{tot}} \cdot P_{tot} \sim 60 \cdot 10^9 \tag{S.28}$$

In the regime where the chromatin is diluted (large amount of metabolites), the NC ratio can be well approximated by $NC_1 = \frac{P_n}{P_c}$. Usual values of NC reported in the literature typically range from 0.3 to 0.6 [7], [8]. We thus estimate reasonable values of $P_n$ as:

$$P_n = \frac{NC}{1 + NC} \cdot P_{tot} \sim 3 \cdot 10^8 \tag{S.29}$$

Note that we also used the numerical solutions of Eq.S.52 in Section VI to infer $NC_1$ from $NC$ exactly. This method made no qualitative difference to the results plotted in Fig.4.

#### K. Estimation of an upper bound for the hydrostatic pressure difference of the nucleus

Even though the stiffness of the lamina layer is susceptible to vary according to the tissue the cell is belonging to [9], its stretching modulus was reported to range from 1 to 25mN/m [10], [11]. Also, Lamina turnover rate is much slower than the actin turnover rate. Together, this suggests that Lamina - at the difference to the cortical actin - can sustain bigger pressure difference on longer timescales. This solid-like behavior of Lamina was observed during micropipette aspiration of Oocyte nuclei through the formation of membrane wrinkles at the pipette entrance [10]. We thus chose to mathematically model lamina with an elastic constitutive equation when it is tensed Eq.S.71. Using Laplace law, we estimate an upper bound for $\Delta P_n$, assuming a typical nuclear radius of 5 $\mu m$, to be:

$$\Delta P_n \sim \frac{2K}{R} \sim 10^4 Pa \tag{S.30}$$

#### L. Estimation of the second virial term in the osmotic pressure

We estimate the steric term in the osmotic pressure to be:

$$\pi_{steric} \sim kT \cdot v_p \cdot p_{tot}^2 \sim 2kPa \tag{S.31}$$

where, $v_p$ is the excluded volume per protein, estimated to be $v_p \sim \frac{R}{P_{tot}} \sim 375nm^3$. This corresponds to a protein radius of 4.5nm, a value coherent with observations [3]. This steric contribution in the osmotic pressure may thus safely be neglected, as $\pi_{steric} << \pi_0$.

### IV. A CELL GROWTH MODEL

We summarize here the equations derived and discussed in the main text (Eqs.6,7). The rates of production of mRNAs and proteins in the non-saturated and saturated regimes read:

$$\dot{M}_j = \begin{cases} k_0 \cdot \phi_j \cdot P_p - \frac{M_j}{\tau_m}, & if \quad P_p \leq P_p^* \\ k_0 \cdot g_j \cdot \mathcal{N}_p^{max} - \frac{M_j}{\tau_m}, & if \quad P_p \geq P_p^* \end{cases} \tag{S.32}$$

$$\dot{P}_j = \begin{cases} k_t \cdot \frac{M_j}{\sum M_j} \cdot P_r - \frac{P_j}{\tau_p}, & if \quad P_r \leq P_r^* \\ k_t \cdot M_j \cdot \mathcal{N}_r^{max} - \frac{P_j}{\tau_p}, & if \quad P_r \geq P_r^* \end{cases} \tag{S.33}$$

The cut-off values - $P_p^*$, $P_r^*$ - above which the substrates become saturated are obtained by imposing continuities of the production rates at the transition:

$$\begin{cases} P_p^* = \mathcal{N}_p^{max} \cdot \sum g_j \\ P_r^* = \mathcal{N}_r^{max} \cdot \sum M_j \end{cases} \tag{S.34}$$

#### A. Neither DNA nor mRNAs are saturated: $P_p \leq P_p^*$ and $P_r \leq P_r^*$

The fast degradation rate of mRNAs ensures that their number reach steady-state during growth (Eq.8 which with Eq.7 yield an exponential growth for the number of ribosomes $P_r = P_{r,0} \cdot e^{k_r \cdot t}$ (with $k_r = k_t \cdot \phi_r - 1/\tau_p$) and of any other protein, $P_j = \frac{\phi_j}{\phi_r} \cdot P_r$; where we neglected the initial conditions on proteins other than ribosomes due to the exponential nature of the growth. Incorporating the dynamics of growth of the enzyme catalyzing the amino-acid biosynthesis $P_e$ into Eq.9, we obtain the number of free amino-acids in the cell:

$$A^f = \left(\phi_e \cdot \frac{k_{cat}}{k_r} - l_p\right) \cdot \frac{P_r}{\phi_r} \tag{S.35}$$

Using the expression of the volume Eq.4 derived from the PLM coupled to our quantitative order of magnitudes, it is straightforward to show that the volume grows exponentially :

$$V = \left( v_p + \frac{(z_{A,f}+1) \cdot \left(\phi_e \cdot \frac{k_{cat}}{k_r} - l_p\right)}{2n_0} \right) \cdot \frac{P_r}{\phi_r} \tag{S.36}$$

where we assumed the dry volume to be mainly accounted by proteins. Incorporating the previous expressions in the equation for the dry mass density Eq.5, we obtain the homeostatic dry mass density written in the main text Eq.10. These expressions were obtained assuming that neither the DNA nor the mRNA were saturated. Importantly, mRNAs cannot be saturated if DNA is not saturated because the cut-off value $P_r^*$ for which ribosomes saturates mRNAs grows at the same speed as the number of ribosomes : $P_r^* = \mathcal{N}_r^{max} \cdot k_0 \cdot \tau_m \cdot \frac{\phi_p}{\phi_r} \cdot P_r$. Hence, DNA will saturate before mRNAs during interphase, at a time $t^*$ given by:

$$t^* = \frac{1}{k_r} \cdot ln \left( \frac{g_r}{g_p} \cdot \frac{\mathcal{N}_p^{max} \cdot \sum g_j}{P_{r,0}} \right) \tag{S.37}$$

#### B. DNA is saturated but not mRNAs: $P_p \geq P_p^*$ and $P_r \leq P_r^*$

The only difference with the previous regime is that mRNA number saturates to the value $M_j = k_0 \cdot g_j \cdot \tau_m \cdot \mathcal{N}_p^{max}$. Hence, the threshold $P_r^*$ will saturate to the value:

$$P_r^* = \mathcal{N}_r^{max} \cdot \mathcal{N}_p^{max} \cdot k_0 \cdot \tau_m \cdot \sum g_j \tag{S.38}$$

This allows for the subsequent saturation of mRNAs by ribosomes after a time $t^{**}$; whose expression can be derived after simple algebra as :

$$t^{**} = t^* + \frac{1}{k_r} \cdot ln \left( \frac{g_p}{g_r} \cdot \mathcal{N}_r^{max} \cdot k_0 \cdot \tau_m \right) \tag{S.39}$$

However, before reaching this time, there won't be any consequence on the proteomic dynamics, which still scales with the number of ribosomes $P_j = \frac{\phi_j}{\phi_r} \cdot P_r$. This regime thus still corresponds to an exponential growth and the dry mass density remains at its homeostatic value Eq.10.

#### C. Both DNA and mRNAs are saturated: $P_p \geq P_p^*$ and $P_r \geq P_r^*$

The dynamics of growth is profoundly impacted by mRNA saturation. The protein number no longer grows exponentially, but saturates to the stationary value $P_j^{stat} = k_t \cdot k_0 \cdot \tau_p \cdot \tau_m \cdot \mathcal{N}_r^{max} \cdot \mathcal{N}_p^{max} \cdot g_j$ after a typical time $t^{**} + \tau_p$ according to:

$$P_j = P_j^{stat} + \left(P_j(t^{**}) - P_j^{stat}\right) \cdot e^{-\frac{t-t^{**}}{\tau_p}} \tag{S.40}$$

The loss of the exponential scaling of proteins implies a breakdown of the proportionality between amino-acid and protein numbers as predicted by the amino-acid biosynthesis equation Eq.9. The total amino-acid pool in the cell $A_{tot} = A_f + l_p \cdot P_{tot}$ now scales as:

$$A_{tot} = A_{tot}(t^{**}) + k_{cat} \cdot \left( P_e^{stat} \cdot (t - t^{**}) - \tau_p \cdot (P_e(t^{**}) - P_e^{stat}) \cdot e^{-(t-t^{**})/\tau_p} \right) \tag{S.41}$$

with, $A_{tot}(t^{**}) = \frac{\phi_e}{\phi_r} \cdot \frac{k_{cat}}{k_r} \cdot \mathcal{N}_r^{max} \cdot \mathcal{N}_p^{max} \cdot k_0 \cdot \tau_m \cdot \sum g_j$. Although, expressions still remain analytical in the transient regime and were implemented in Fig.2 in order to quantitatively test our theory, we avoid analytical complications here, by writing expressions after saturation has been reached, i.e., after a typical time $t^{**} + \tau_p$. The volume thus increases linearly with time:

$$V^{lin} = v_p \cdot P_{tot}^{stat} + \frac{(z_{A^f} + 1) \cdot (A_{tot}(t^{**}) + k_{cat} \cdot P_e^{stat} \cdot (t - t^{**}) - l_p \cdot P_{tot}^{stat})}{2n_0} \tag{S.42}$$

As emphasized in the main text, the fundamental property of this regime is that the dry mass density is predicted to decrease with time with no other mechanism than a simple crowding effect on mRNAs (see Eq.11 in the main text).

#### D. Quantification of the model of growth with published data

Many of the parameters involved in the growth model can be obtained independently, so that four parameters suffice to fully determine the volume, the amount of protein and the dry mass density during interphase growth. Here, we summarize the equations used to fit the data displayed in Fig.2. The volume can be expressed as:

$$V = \begin{cases} v_1 \cdot e^{k_r \cdot t}, & if \quad t \leq t^{**} \\ v_2 \cdot (t - t^{**}) + v_3 \cdot e^{-(t-t^{**})/\tau_p} + v_4, & if \quad t \geq t^{**} \end{cases} \tag{S.43}$$

in which $(v_1,\ v_2,\ v_3,\ v_4)$ are volumes that can be, if needed, expressed function of the previously defined parameters. We obtain $(v_2,\ v_3,\ v_4)$ as a function of $v_1,\ \tau_p$ and $t^{**}$ by imposing regularity constraints on the volume and growth rate:

$$\begin{cases} v_3 = \tau_p^2 \cdot k_r^2 \cdot v_1 \cdot e^{k_r \cdot t^{**}} \\ v_2 = \frac{v3}{\tau_p} + k_r \cdot v_1 \cdot e^{k_r \cdot t^{**}} \\ v_4 = v_1 \cdot e^{k_r \cdot t^{**}} - v_3 \end{cases} \tag{S.44}$$

Similarly, the normalized total number of protein can be expressed as:

$$\frac{P_{tot}}{P_{tot}(1h)} = \begin{cases} e^{k_r \cdot (t-1)}, & if \quad t \leq t^{**} \\ p_1 + p_2 \cdot e^{-(t-t^{**})/\tau_p}, & if \quad t \geq t^{**} \end{cases} \tag{S.45}$$

Again imposing regularity constraints at the mRNA saturating transition, allows us to relate $p_1$ and $p_2$ to $k_r$, $\tau_p$ and $t^{**}$.

$$\begin{cases} p_2 = -\tau_p \cdot k_r \cdot e^{k_r \cdot (t^{**}-1)} \\ p_1 = e^{k_r \cdot (t^{**}-1)} - p_2 \end{cases} \tag{S.46}$$

Finally, we can express the buoyant mass density $\rho^b$ of the cell (see Fig.2 for a definition) using the expressions of total protein number and volume Eq.S.45,S.43 :

$$\frac{\rho^b - \rho^w}{\rho^{b,0} - \rho^w} = \begin{cases} 1, & if \quad t \leq t^{**} \\ \frac{P_{tot}}{P_{tot}(0h)} \cdot \frac{v_1}{V}, & if \quad t \geq t^{**} \end{cases} \tag{S.47}$$

We use a density of water 4% larger than that of pure water ($\rho^{w,eff} = 1.04$kg/L instead of $\sim 1$kg/L) to compensate for our approximation to consider the dry mass as entirely made of proteins. Proteins are known to only occupy $\%_p^{mass} = 0.6$ of the dry mass, itself being of order $\rho = 0.1$kg/L Tab.S1. Thus, we simply use as the effective water mass density, $\rho^{w,eff} = \rho^w + (1 - \%_p^{mass}) \cdot \rho \sim 1.04$kg/L.

#### E. Fitting procedure

We detail in this appendix the method used to determine the four fitting parameters: $\tau_p$, $t^{**}$, $k_r$, $v_1$ from the cell volume data Fig.2.B. Our model (Eq.S.43,S.44) displays two different regimes of growth according to the saturation state of mRNAs. Our fitting procedure is thus divided into two steps. First, we impose an arbitrary transition time $t^{**}$ to determine by a least mean square minimization the three other parameters. Then, we minimize the variance between the obtained solution with the data to determine $t^{**}$. The optimal values of the fitting parameters are:

$$t^{**} = 2h44\text{min} \quad , \quad \tau_p = 1h9\text{min} \quad , \quad k_r = 0.62\text{h}^{-1} \quad , \quad v_1 = 30\text{fL} \tag{S.48}$$

### V. MANNING CONDENSATION

We give a simple description of the phenomenon of Manning condensation, based on [12]. The electrostatic potential close to an infinitely charged thin rod, in a salt bath, reads:

$$\psi = \frac{2 \cdot l_b}{A} \cdot ln(\kappa r) \tag{S.49}$$

where, $l_b$ is the Bjerrum length. It is the length at which the elecrostatic interaction between two elementary particles is on the order of kT. Its value in water at room temperature is $l_B \approx 0.7$nm. A is the average distance between two charges on the polymer. $\kappa^2 = 8\pi l_B(n^+ + n^-)$ is the inverse of the Debye length. At equilibrium, the distribution of charges around the rod follows a Boltzmann distribution:

$$n^+ = n_0 \cdot e^{-\Psi} = \frac{n_0}{(\kappa r)^{2 \cdot \frac{l_b}{A}}} \tag{S.50}$$

The total number of positive charges per unit length of the rod reads within a distance $\mathcal{R}$:

$$N(\mathcal{R}) = \int_0^{\mathcal{R}} n^+ 2\pi r dr = \frac{2\pi \cdot n_0}{\kappa^{2 \cdot \frac{l_b}{A}}} \cdot \int_0^{\mathcal{R}} \frac{1}{(r)^{2\frac{l_b}{A}-1}} dr \tag{S.51}$$

When $u = \frac{l_B}{A} < 1$, $N(\mathcal{R})$ is dominated by its upper bond and goes to 0 close to the rod. On the other hand, when $u = \frac{l_B}{A} > 1$, $N(\mathcal{R})$ diverges as $\mathcal{R} \to 0$ indicating a strong condensation of the counterions on the rod. This singularity is symptomatic of the breakdown of the linear Debye-Huckel theory. The solution of the nonlinear Poisson-Boltzmann equation shows that there is formation of a tighly bound layer of counterions very near the rod, which effectively decreases the charge density (increases $A$) up to the value $u_{eff} = 1$ [13]. It means that if $A$ is smaller than $l_B$ the manning condensation will renormalize $A$ to $A^{eff} = l_B$. The rationale behind this renormalization is to decrease the electrostatic energy of the system by condensing free ions on the polymer. Note that there is an energy penalty associated to the loss of entropy of the condensed counterions. For weakly charged polymers this loss of entropy is not energetically favorable - case where $u = \frac{l_B}{A} < 1$ and no condensation occurs. If the density of charge of the polymer increases, Manning condensation becomes energetically favorable - case where $u = \frac{l_B}{A} > 1$. By virtue of the high lineic charge of DNA, Manning condensation will be favorable, $u^{DNA} \sim 4$.

### VI. THE NESTED PLM MODEL

The nested PLM is described by a set of non-linear equations, i.e., the electroneutrality, the balance of pressures, and the balance of ionic fluxes, in the cytoplasm, subscript c, and in the nucleus, subscript n. In its most general form the system reads:

$$\begin{cases} n_c^+ - n_c^- - z_p \cdot p_c - z_a \cdot a_c = 0 \\ n_n^+ - n_n^- - z_p \cdot p_n - z_a \cdot a_n - q = 0 \\ \Delta\Pi_c = \Delta P_c \\ \Delta\Pi_n = \Delta P_n \\ n_c^+ \cdot n_c^- = \alpha_0 \cdot n_0^2 \\ n_n^+ \cdot n_n^- = \alpha_0 \cdot n_0^2 \\ (n_n^+)^{z_a} \cdot a_n = (n_c^+)^{z_a} \cdot a_c \end{cases} \tag{S.52}$$

Here, we apply the nested PLM to mammalian cells, such that we can neglect the cytoplasmic difference of hydrostatic pressure with respect to the external osmotic pressure. If the NE is not under tension, the condition of osmotic balance at the NE simply implies that the volume of each compartment takes the same functional form as in the PLM model:

$$\begin{cases} V_n = R_n + \frac{N_n^{tot}}{2n_0} \\ V_c = R_c + \frac{N_c^{tot}}{2n_0} \end{cases} \tag{S.53}$$

It is thus straightforward to show that both the volume of the nucleus and the volume of the cytoplasm scale with each other Eq.12.

#### A. Dry volumes in the nucleus and in the cytoplasm

We assume that the dry volumes in the nucleus and in the cytoplasm are proportional to the total volumes of each compartments and are equal to each other: $R_n = r \cdot V_n$ and $R_c = r \cdot V_c$. Under this assumption the NC ratio simply becomes the ratio of the wet volumes:

$$NC = \frac{V_n}{V_c} = \frac{V_n - R_n}{V_c - R_c} \tag{S.54}$$

This hypothesis is practical rather than purely rigorous. It is based on experiments that suggest that dry mass occupies about 30% of the volume of both the nucleus and the cytoplasm for several cell types and conditions [14], [15],[16]. Nonetheless, even if this assumption were to be inexact, our discussion would then rigorously describe the slope of the linear relationship between nucleus and cell volume (Eq.12) which was shown to be robust to perturbation [7].

#### B. Membrane potential in the simple PLM model

Using the results provided earlier we find that a transmembrane potential exists as soon as there are trapped charged particles. The plasma membrane potential reads:

$$\begin{cases} U = ln\left(\frac{-z\cdot(-1+r)+\sqrt{z^4+\alpha_0-z^2\cdot(-1+\alpha_0+2\cdot r)}}{z^2-1}\right) \\ \text{With,} \quad r = \sqrt{1+(z^2-1)\cdot(1-\alpha_0)} \end{cases} \tag{S.55}$$

We find that $U$ monotonically increases (in absolute value) with the average charge of the cell trapped components. This differs from the nuclear membrane potential that vanishes when the charge of the chromatin is diluted regardless of the properties of the trapped proteins Eq.14.

#### C. General Formula for the regime $NC_2$, i.e., no metabolites

As stated in the main text an important limit regime, $NC_2$, is achieved when there are no metabolites in the cell. Specifically, the previous system of equations becomes uncoupled with respect to the nuclear and cytoplasmic set of variables such that we can solve the system analytically. Using the exact same algebra as used in the simple PLM we express the volumes and the NC ratio as:

$$\begin{cases} V_{tot} = (R_c + R_n) + \frac{N_n^{tot}+N_c^{tot}}{2n_0} \\ N_c^{tot} = P_c \cdot \frac{z_p^2-1}{-1+\sqrt{1+(1-\alpha_0)\cdot(z_p^2-1)}} \\ N_n^{tot} = P_n \cdot \frac{(z_{n,eff}^2-1)}{-1+\sqrt{1+(1-\alpha_0)(z_{n,eff}^2-1)}} \\ z_{n,eff} = z_p + \frac{Q^{eff}}{P_n} \\ NC_2 = NC_1 \cdot \frac{(z_{n,eff}^2-1)}{(z_p^2-1)} \cdot \frac{-1+\sqrt{1+(1-\alpha_0)(z_p^2-1)}}{-1+\sqrt{1+(1-\alpha_0)(z_{n,eff}^2-1)}} \end{cases} \tag{S.56}$$

#### D. Analytical solutions in the regime $z_p = 1,\ z_a = 1$, and $\alpha_0 \sim 0$

In this regime of high pumping, no anions occupy the cell. We simplify the notations by denoting by $n$ the concentration of cations. The system of equation to solve is stated in the main text Eq.13. We first express the concentrations of cations and metabolites in the cytoplasm and nucleus as a function of $n_0$, $q$, and $p$ thanks to the electroneutrality equations and balance of osmotic pressures:

$$\begin{cases} n_n = n_0 + \frac{q}{2} \\ a_n^f = \left(n_0 - \frac{q}{2}\right) - p_n \\ n_c = n_0 \\ a_c^f = n_0 - p_c \end{cases} \tag{S.57}$$

This allows us to write the NE potential as Eq.14 in the main text. Using the balance of nuclear osmotic pressure we express the nuclear volume function of the number of nuclear osmolytes:

$$V_n - R_n = \frac{Q^{eff} + 2A_n^f + 2P_n}{2n_0} \tag{S.58}$$

This implies that the NE potential can be written without the dependence on $n_0$ as in Eq.14. We then express the NC ratio in two different manners. First, using the interpretation of wet volumes namely, the total number of osmolytes in the compartments over $2n_0$. Second, we take advantage of the concentrations of metabolites and cations in Eq.S.57 to express the ratio of protein concentrations. After simple algebra we obtain:

$$\begin{cases} NC = \frac{P_n}{P_c} \cdot \frac{p_c}{p_n} = NC_1 \cdot \left(1 + \frac{Q^{eff}}{2P_n + 2A_n^f + Q^{eff}} + \frac{{Q^{eff}}^2}{2P_n + 2A_n^f + Q^{eff}}\right) \\ NC = \frac{1}{2} \cdot \frac{2A_n^f + 2P_n + Q^{eff}}{A_{tot}^f - A_n^f + P_c} \end{cases} \tag{S.59}$$

For clarity, we now normalize each number by $2P_n$, e.g, $\overline{A}_{tot} = \frac{Atot}{2P_n}$. Equating both expressions of the NC ratio leads to a second order polynomial in $\overline{A}_n$ :

$$2(1 + \frac{1}{NC_1}) \cdot \overline{A}_n^2 + \left(-2\overline{A}_{tot} + (1 + \overline{Q}^{eff})^2 + \frac{1}{NC_1} \cdot (1 + 2\overline{Q}^{eff})\right) \cdot \overline{A}_n - \overline{A}_{tot} \cdot (1 + \overline{Q}^{eff})^2 = 0 \tag{S.60}$$

The solution $\overline{A}_n$ now reads:

$$\overline{A}_n = \frac{2\overline{A}_{tot} - \frac{1}{NC_1} \cdot (1 + 2\overline{Q}^{eff}) - (1 + \overline{Q}^{eff})^2 + \sqrt{\left(2\overline{A}_{tot} - \frac{1}{NC_1} \cdot (1 + 2\overline{Q}^{eff}) - (1 + \overline{Q}^{eff})^2\right)^2 + 8 \cdot (1 + \frac{1}{NC_1}) \cdot \overline{A}_{tot} \cdot (1 + \overline{Q}^{eff})^2}}{4 \cdot (1 + \frac{1}{NC_1})} \tag{S.61}$$

Which leads to the following expression for NC:

$$NC = NC_1 \cdot \frac{2\overline{A}_{tot} + \frac{1}{NC_1} + (1 + \overline{Q}^{eff})^2 + \sqrt{\left(2\overline{A}_{tot} - \frac{1}{NC_1} \cdot (1 + 2\overline{Q}^{eff}) - (1 + \overline{Q}^{eff})^2\right)^2 + 8 \cdot (1 + \frac{1}{NC_1}) \cdot \overline{A}_{tot} \cdot (1 + \overline{Q}^{eff})^2}}{2 \cdot \left(1 + 2\overline{A}_{tot} + \frac{1}{NC_1} + \overline{Q}^{eff}\right)} \tag{S.62}$$

As a sanity check, we verify some asymptotic expressions discussed in the main text. For example, when $\overline{Q}^{eff} << 1$ or $\overline{A}_{tot} >> 1$ we recover that NC becomes equal to $NC_1$. On the other hand, when $\overline{A}_{tot} << 1$, we recover that $NC = NC_1 \cdot (1 + \overline{Q}^{eff}) = NC_2$

#### E. Control parameters of the nested PLM during growth

The precise value of the parameter $NC_1 = \frac{P_n}{P_c}$ is biologically set by an ensemble of complex active processes ranging from transcription, translation to the Ran GTPase cycle and nuclear transport. The precise modelling of nucleo-cytoplasmic transport is out of the scope of this paper but could easily be incorporated to our framework. Nonetheless, we can safely assume that nucleo-cytoplasmic transport is fast compared to the typical timescale of growth. In this case, neglecting protein degradation on the timescale of the G1 phase, the total number of proteins in the nucleus is simply the number of proteins assembled that possessed a nuclear import signal (NIS) in their sequence. Using the same notation as earlier, in the exponential growth regime, the total number of proteins in the nucleus reads:

$$P_{tot,n}(t) = \sum_{j \in NIS} \frac{\phi_j}{\phi_r} \cdot P_r(t) \tag{S.63}$$

where $P_r(t)$ accounts for the number of ribosomes, $\phi_j$ is the fraction of genes coding for the protein j (see Appendix IV). The subscript j is summed over the genes coding for proteins having nuclear import signals in their sequence. Proteins in the nucleus can either be DNA bound or unbound. For example, histones or DNA polymerases bind to the DNA. Only the unbound proteins contribute to the osmotic pressure. Denoting $k_{u,j}$ and $k_{b,j}$ the reaction rate of binding and unbinding of protein j and assuming that the reactions of binding and unbinding are fast compared to the timescale of growth, we finally express the number of free proteins in the nucleus as :

$$P_{free,n}(t) = \sum_{j \in NIS} \frac{k_{u,j}}{k_{b,j} + k_{u,j}} \cdot \frac{\phi_j}{\phi_r} \cdot P_r(t) \tag{S.64}$$

It is then straightforward to express $NC_1$ as:

$$NC_1 = \frac{\sum_{j \in NIS} \frac{k_{u,j}}{k_{b,j}+k_{u,j}} \cdot \frac{\phi_j}{\phi_r}}{\sum_{j \notin NIS} \frac{k_{u,j}}{k_{b,j}+k_{u,j}} \cdot \frac{\phi_j}{\phi_r}} \tag{S.65}$$

An important result of this abstract modelling is that $NC_1$ is independent of time during the exponential growth due to the fact that both $P_n(t)$ and $P_c(t)$ are proportional to $P_r(t)$, which is why we adopted it as a control parameter. The same goes for our second control parameter $\frac{A_{tot}}{2P_n}$, which is also constant during exponential growth.

#### F. Phase Diagram

In this paragraph we address the case $\Delta P_n \neq 0$ and assume that the cell does not adhere to the substrate such that we consider the nucleus to be spherical. For simplicity, we neglect the dry volume because we want to consider hypo-osmotic shock experiment where dry mass will be diluted, making a dry volume a second order effect of the order 10%. We first make the problem dimensionless. There are two dimensions in our problem: an energy and a length. This means that we can express all our parameters that possess a dimension with a unit energy and a unit length. Moreover, we have three parameters with physical dimensions: the extracellular osmolarity $n_0$, the NE tension $\gamma$, and the thermal energy $kT$. The theorem of Buckingham [17] tells us that we can fully describe our problem with a single dimensionless parameter and the 3 parameters with by definition no dimensions. In this geometry, we choose, $[\left(\frac{4\pi}{3}\right)^{1/3} \cdot \frac{\gamma_0}{kTX_n^{1/3} n_0^{2/3}}, \alpha_0, z_n, X_n]$. Laplace law reads:

$$\Delta P_n = \frac{2\gamma_n}{(\frac{3}{4\pi}V_n)^{1/3}} \tag{S.66}$$

Using the following dimensionless quantities:

$$\begin{cases} \overline{V}_n = \frac{2n0}{X_n} \cdot V_n \\ \overline{\gamma}_n = \left(\frac{4\pi}{3}\right)^{1/3} \cdot \frac{\gamma_0}{kTX_n^{1/3} n_0^{2/3}} \end{cases} \tag{S.67}$$

Equality of pressures becomes:

$$\sqrt{\left(\frac{z_n}{\overline{V}_n}\right)^2 + \alpha_0} + \frac{1}{\overline{V}_n} - 1 - \frac{\overline{\gamma}_n}{\overline{V}_n^{1/3}} = 0 \tag{S.68}$$

Eq.S.68 cannot be solved analytically for $\overline{V}_n$. However, five asymptotic regimes can be identified (see Fig.S1):

$$\begin{cases} V_1 = \left(\frac{3}{16\pi}\right)^{\frac{1}{2}} \cdot \left(\frac{kTX_n}{z_n \cdot \gamma_n}\right)^{\frac{3}{2}} \\ V_2 = \left(\frac{3}{2^{10}\cdot\pi}\right)^{\frac{1}{5}} \cdot \left(\frac{kT\cdot z_n^2 \cdot X_n^2}{n_0\cdot\sqrt{\alpha_0}\cdot\gamma_n}\right)^{\frac{3}{5}} \\ V_3 = \left(\frac{3}{16\pi}\right)^{\frac{1}{2}} \cdot \left(\frac{kT\cdot X_n}{\gamma_n}\right)^{\frac{3}{2}} \\ V_4 = \frac{X_n}{2n_0} \cdot \frac{1}{1-\sqrt{\alpha_0}} \\ V_5 = \frac{X_n}{2n_0} \cdot \frac{z_n}{\sqrt{1-\alpha_0}} \end{cases} \tag{S.69}$$

- $V_4$ and $V_5$ are the limit regimes where osmotic pressure is balanced at the NE.
- $V_3$ is the limit regime where the difference of osmotic pressure is dominated by the impermeant molecules trapped inside the nucleus. This happens when the proteins are not or very weakly charged. This difference of osmotic pressure is balanced by the Laplace pressure of the lamina.
- $V_1$ is the limit regime where the difference of osmotic pressure is dominated by the counterions of the impermeant molecules. This difference of osmotic pressure is balanced by the Laplace pressure of the lamina.
- $V_2$ is an intermediate regime that can arise when $\alpha_0 \approx 1$. The difference of osmotic pressure takes the form of $\Delta\Pi_n \approx kT \cdot \frac{1}{\sqrt{\alpha_0}} \cdot \frac{(z_n \cdot x_n)^2}{4n_0}$. This osmotic pressure defines an effective virial coefficient between monomers of DNA and proteins $v_{el} = \frac{1}{\sqrt{\alpha_0}} \frac{z_n^2}{2n_0}$. This difference of osmotic pressure is balanced by the Laplace pressure at the NE.
- Note that when $\alpha_0 \approx 0$ (strong pumping), only the counterion necessary for electroneutrality remain in the nucleus. $\Pi_n$ is simply $(z_n + 1) \cdot x_n$ and is either balanced by the Laplace pressure of the lamina or the external osmotic pressure (see Fig.S1)

Finally, the analytical expressions for the crossover lines $\overline{\gamma}_{i,j}$ between regime of volume $V_i$ and volume $V_j$, plotted in Fig.S1 read :

$$\begin{cases} \overline{\gamma}_{1,2} = (4 \cdot \alpha_0 \cdot z_n)^{\frac{1}{3}} \\ \overline{\gamma}_{1,4} = z_n \cdot (1 - \sqrt{\alpha_0})^{\frac{2}{3}} \\ \overline{\gamma}_{1,5} = z_n^{\frac{1}{3}} \cdot (1 - \alpha_0)^{\frac{1}{3}} \\ \overline{\gamma}_{2,5} = \frac{z_n^{\frac{1}{3}} \cdot (1-\alpha_0)^{\frac{5}{6}}}{2\cdot\sqrt{\alpha_0}} \\ \overline{\gamma}_{2,4} = \frac{z_n^2 \cdot (1-\sqrt{\alpha_0})^{\frac{5}{3}}}{2\cdot\sqrt{\alpha_0}} \\ \overline{\gamma}_{2,3} = z_n^{-\frac{4}{3}} \cdot (4 \cdot \alpha_0)^{\frac{1}{3}} \\ \overline{\gamma}_{3,5} = z_n^{-\frac{2}{3}} \cdot (1 - \alpha_0)^{\frac{1}{3}} \\ \overline{\gamma}_{3,4} = (1 - \sqrt{\alpha_0})^{\frac{2}{3}} \\ z_{1,3} = 1 \\ z_{4,5} = \frac{\sqrt{1-\alpha_0}}{1-\sqrt{\alpha_0}} \end{cases} \tag{S.70}$$

#### G. Saturating volume after a hypo-osmotic shock

The saturation occurs when the nuclear osmotic pressure is balanced by the Laplace pressure making nuclear volume insensitive to the external osmolarity Eq.S.69. We assume that the NE behaves elastically with a stretching modulus $K$ beyond a surface area $S^*$ for which NE folds are flattened:

$$\gamma_n = \begin{cases} 0 & , if \quad S_n \leq S^* \\ K \cdot \left(\frac{S_n}{S^*} - 1\right) & , if \quad S_n \geq S^* \end{cases} \tag{S.71}$$

As justified in the main text, metabolites tend to leave the nucleus with decreasing external osmolarity. The saturating volume is obtained when $\Delta P_n >> \pi_0$ and $A_n^f << P_n$. From Eq.S.12 applied to the volume of the nucleus, we thus obtain :

$$\Delta P = (z_n^{eff} + 1) \cdot \frac{P_n}{V_n^{max}} \tag{S.72}$$

where, $z_n^{eff} = z_p + \frac{Q}{P_n}$. Similarly to the last subsection, we normalize tensions by $(\frac{3}{4\pi})^{1/3} \cdot kT \cdot P_n^{1/3} \cdot n_0^{2/3}$ and volumes by $\frac{2 \cdot P_n}{2n_0}$. Eq.S.72 leads to the equation ruling the saturating volume :

$$(v_n^{max})^{4/3} - (v_n^{iso})^{2/3} \cdot (1+s) \cdot (\frac{n_0}{n_0^{iso}})^{2/3} \cdot (v_n^{max})^{2/3} - (v_n^{iso})^{2/3} \cdot (1+s) \cdot (\frac{n_0}{n_0^{iso}})^{2/3} \cdot \frac{\frac{1}{2} \cdot (z_n^{eff}+1)}{\overline{K}} = 0 \tag{S.73}$$

where, $s = \frac{S^*}{S_n^{iso}} - 1$ is the fraction of folds that the nucleus possesses at the isotonic osmolarity. $v_n^{max} = \frac{2n_0 \cdot V_n^{max}}{2P_n}$ is the normalized saturating nuclear volume and $v_n^{iso}$ the normalized nuclear volume at the isotonic osmolarity. $\overline{K} = \frac{K}{(\frac{3}{4\pi})^{1/3} \cdot kT \cdot P_n^{1/3} \cdot n_0^{2/3}}$ is the normalized effective stretching modulus of the NE. Solving the previous equation, coming back to real volumes $\frac{V_n^{max}}{V_n^{iso}} = \frac{v_n^{max}}{v_n^{iso}} \cdot \frac{n_0^{iso}}{n_0}$ and taylor develop the result for $n_0 \longrightarrow 0$ leads to Eq.17 in the main text.

#### H. Geometrical impact

The previous equations were conducted for a spherical geometry. Interestingly, while the precise geometry does not qualitatively change our results, we expect the saturation of nuclear volume to occur more easily for a pancake shape - a shape closer to the shape of adhered cells. Indeed, the scaling between surface and volume is approximatively linear in this case: $V \sim h \cdot S$, while it is sub-linear for spheres $S \sim V^{2/3}$. Thus, smaller osmotic shocks will be required to tense the NE and so as to reach the saturating regime.

### VII. ELECTROSTATIC INTERACTIONS ARE ENCOMPASSED WITHIN OUR FRAMEWORK

We directly compute the contribution of electrostatic interactions to the osmotic pressure based on [12]. The total interaction energy of a solution of charged particles of average density $x$ within a volume $V$ is, using the Poisson-Boltzmann framework:

$$\frac{E_{el}}{kT} = \frac{l_B \cdot z^2}{2} \int\int x(\vec{r}) \cdot x(\vec{r'}) \cdot \frac{e^{-\kappa|\vec{r}-\vec{r'}|}}{|\vec{r}-\vec{r'}|} \quad d^3\vec{r} \cdot d^3\vec{r'} \tag{S.74}$$

where $x(\vec{r})$ is the local density of impermeant molecules in the cell. Fourier analysis allows us to rewrite this equation:

$$\frac{E_{el}}{kT} = \frac{l_B \cdot z^2}{2} \int x(\vec{k}) \cdot x(-\vec{k}) \cdot \frac{4\pi}{k^2+\kappa^2} \quad d^3\vec{k} \approx \frac{l_B \cdot z^2}{2} \cdot x^2 \cdot \frac{4\pi}{\kappa^2} \cdot V \tag{S.75}$$

From which we derive the expression of the osmotic pressure:

$$\frac{\pi_{el}}{kT} \approx \frac{1}{2} \cdot \frac{z^2}{2n_0} \cdot x^2 \tag{S.76}$$

We now show that this term is already encompassed within our framework. For the simplicity of the discussion we neglect pumping, i.e., $\alpha_0 \sim 1$. The difference of osmotic pressure then reads (see Eq.S.8):

$$\frac{\Delta\pi}{kT} = \sqrt{(zx)^2 + 4n_0^2} + x - 2n_0 \tag{S.77}$$

which, under the right regime, i.e., $zx << 2n_0$, leads to the same term. As mentioned above, this osmotic pressure defines an effective electrostatic virial coefficient between monomers:

$$v_{el} = \frac{z^2}{2n_0} \tag{S.78}$$

### VIII. POSSIBLE EXTENSION TO EXPLAIN THE SCALING OF OTHER ORGANELLES

Organelles are also known to display characteristic scaling trends with cell size ([18]). Eventhough these scalings may be of different origins and would require much careful treatment with respect to the specificity of the organelle, we highlight in this subsection that our model can easily be extended to also include organelles.
We model an organelle in our theory by a compartment bound by a membrane that trap some molecules. For the sake of generality we assume that there is an active transport of cations through this membrane. As a matter of coherence with the previous notations we will call by $\alpha_{org} = e^{-\frac{p_{org}}{g+}}$ the parameter that compares the active pumping through the organelle's membrane versus the passive leakage. Donnan Equilibrium on both side of the organelle reads :

$$n^+_{org} \cdot n^-_{org} = \alpha_{org} \cdot (n^+_c \cdot n^-_c) = \alpha_{org} \cdot \alpha_0 \cdot n_0^2 \tag{S.79}$$

Hence, the results derived previously also apply to the organelle provided the parameter $\alpha_0$ is changed into $\alpha_{org} \cdot \alpha_0$. Interestingly, in the case of osmotic balance at the membrane of the organelle, it is straightforward to show that the the volume of the organelle also scales with the cell volume:

$$\begin{cases} V_{org} = \left(\frac{N^{tot}_{org}}{N^{tot}}\right) \cdot V_{tot} + \left[\left(\frac{N^{tot}_c + N^{tot}_n}{N^{tot}}\right) \cdot R_{org} - \left(\frac{N^{tot}_{org}}{N^{tot}}\right) \cdot (R_c + R_n)\right] \\ N^{tot} = N^{tot}_c + N^{tot}_n + N^{tot}_{org} \\ N^{tot}_{org} = X_{org} \cdot \frac{(z^2_{org} - 1)}{-1 + \sqrt{1 + (1 - \alpha_0 \cdot \alpha_{org})(z^2_{org} - 1)}} \end{cases} \tag{S.80}$$

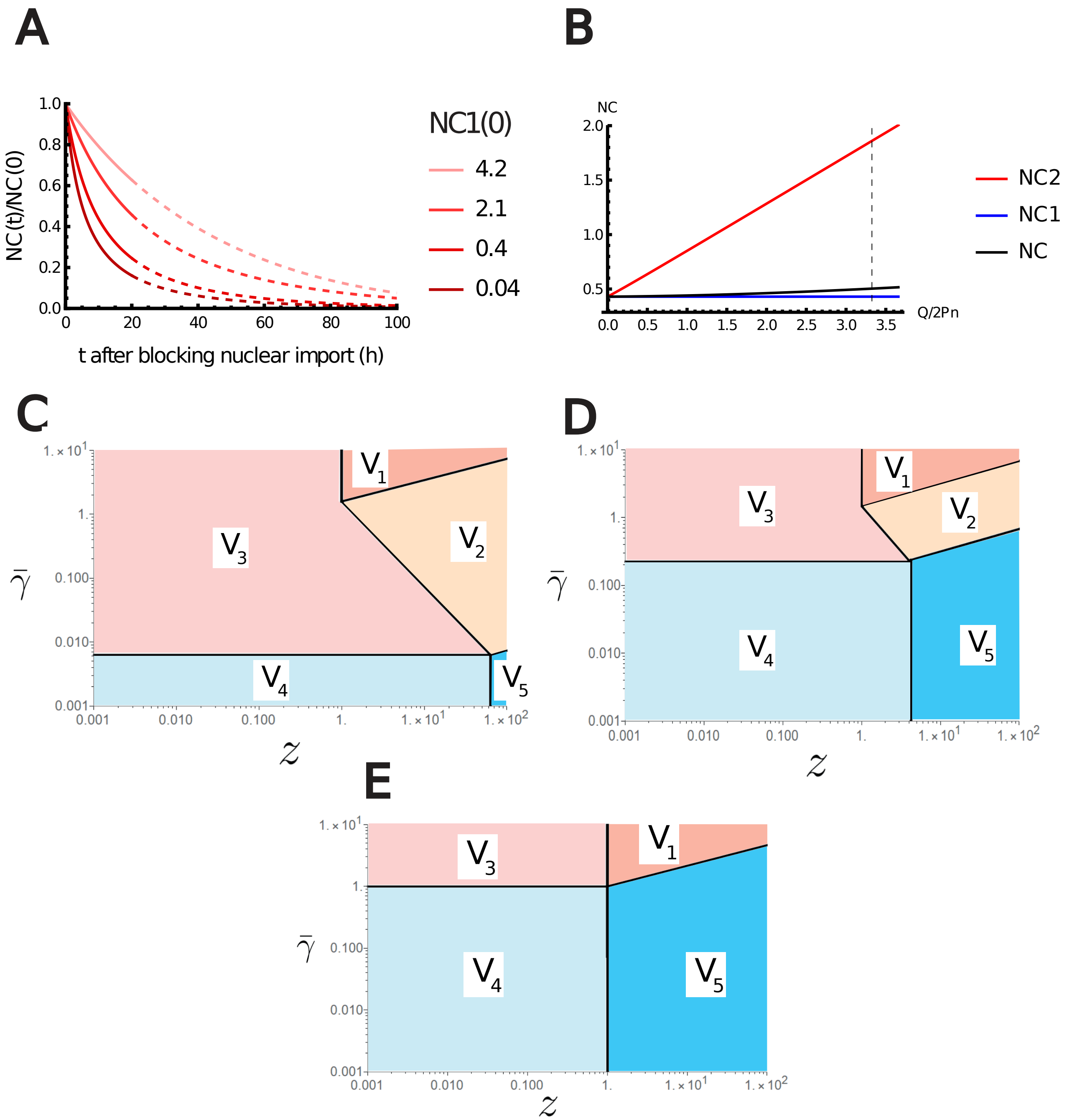


FIG. S1. Additional results of the Nested PLM. (A) Variation of the NC ratio during growth after blocking nuclear import. (B) Variations of the NC ratio according to the effective charge of the chromatin normalized by the number of trapped proteins in the nucleus $\frac{Q}{2P_n}$. The NC ratio is bounded by two limit regimes. NC1, if the number of metabolites is assumed infinite. NC2, if there are no metabolites. The vertical black dashed line depicts the value of $\frac{Q}{2P_n}$ estimated in Appendix III for diploid mammalian cells (C) to (E) Log-Log plot of the different regimes of V Eq.S.69 in the plan $(z_n, \overline{\gamma}_n)$ for $\alpha_0$ fixed (C) $\alpha_0 = 0.99$ (D) $\alpha_0 = 0.8$ (E) $\alpha_0 = 0.001$. The crossover lines plotted are given in Eq.S.70

TABLE S1. Description and values of the parameters used for the order of magnitudes.

| Symbol | Typical Value | Meaning |
|---|---|---|
| $\rho$ | $0.1\text{kg.L}^{-1}$ | Typical dry mass density in a mammalian cell [2] |
| $\mathcal{M}_a$ | 100Da | Average mass of an amino-acid [3] |
| $l_p$ | 400a.a | Average length of an eukaryotic protein [3] |
| $\mathcal{M}_{mRNA}$ | $3 \cdot \mathcal{M}_a$ | Average mass of a mRNA [3] |
| $l_{mRNA}$ | $3 \cdot l_p$ | Average length of a mRNA [3] |
| $l_{bp}$ | 1/3nm | Average length of one base pair |
| $Q_{bp}$ | 2 | Average number of negative charge per base pair |
| $L_{nucleosome}$ | 200bp | Average length of DNA per nucleosome |
| $L_{link}$ | 53bp | Length of the DNA linking two histones |
| $L_{wrap}$ | 147bp | Length of the DNA wrapped around one histone |
| $u_{DNA}$ | 4 | Manning parameter for pure DNA, i.e., 75% of the charges will be screened by manning condensation. |
| $L_{tot}$ | $6 \cdot 10^9$bp | Total length of the DNA within a diploid human cell |
| $Q_{hist}$ | 76 | Average number of positive charges per histone at less than 1nm from the wrapped DNA backbone [5] |
| $Q_{wrap}$ | 174 | Average number of condensed counterions around the wrapped DNA [5] |
| $l_b$ | 0.7nm | Bjerrum length in water at 300k |
| K | 25mN/m | Stretching modulus of Lamina [10] |

[1] O. Sten-Knudsen, *Biological Membranes: Theory of Transport, Potentials and Electric Impulses* (Cambridge University Press, 2007).

[2] E. Zlotek-Zlotkiewicz, S. Monnier, G. Cappello, M. Le Berre, and M. Piel, Journal of Cell Biology **211**, 765 (2015), https://rupress.org/jcb/article-pdf/211/4/765/1370799/jcb_201505056.pdf.

[3] R. Philipps and R. Milo, *Cell Biology by the numbers* (CRC Press, 2015).

[13] H. Schiessel, *Biophysics for beginners: a journey through the cell nucleus* (Pan Stanford Pub., 2014).
